# Transcriptional signature pattern in black, blue and purple wheat and impact on seed pigmentation and other associated features: Comparative transcriptomics, genomics and metabolite profiling

**DOI:** 10.1101/2022.05.21.492912

**Authors:** Saloni Sharma, Ashish Kumar, Dalwinder Singh, Anita Kumari, Payal Kapoor, Satveer Kaur, Bhawna Sheoran, Monika Garg

**Affiliations:** National Agri-Food Biotechnology Institute, Mohali, Punjab, 140306-India; International Centre for Genetic Engineering and Biotechnology, New Delhi, 110067-India

**Keywords:** anthocyanin, wheat, stress, transcriptomics, transcription factor

## Abstract

Anthocyanin biosynthesis in plants is complex, especially in a polyploid monocot wheat plant. Using whole-genome sequencing, transcriptomics, and LC-MS/MS, we investigated anthocyanin production in pigmented (black, blue, and purple) wheat seeds. According to differential gene expression profiling, 2AS-MYC, 7DL-MYB, WD40 regulatory genes controls purple pericarp coloration, 4DL-MYC, 2AS-MYC, 7DL-MYB, WD40 controls blue aleurone coloration, and 4DL-MYC, 7DL-MYB, WD40 controls black aleurone colour. We believe that at least one MYC and MYB isoform is sufficient to regulate the anthocyanin synthesis in pericarp or aleurone. Based upon the reduced expressions of the genes belonging to the 4D, SSR molecular marker mapping, variant calling using genome sequencing and IGV browser gene structure visualization, it was inferred that the advanced black and blue wheat lines were substitution lines (4E{4D}), with very small recombinations. Pericarp anthocyanin profiling is controlled by a mutation in chromosome 2AS of purple wheat, and environmental variations more influence pigmented pericarp trait. The expression patterns of anthocyanin structural and other genes varied in different colored wheat, corroborating differences in agronomical metrics.

## Introduction

Cereals, particularly whole grains, make up a large part of the global food supply and play an important role in maintaining a healthy lifestyle. As a result, numerous researchers and organisations continue to explore the potential of whole grains, focusing more on bioactive components that are suited for developing high-value-added food items with enhanced health benefits. The notion of a healthy diet has become so widespread that individuals are becoming increasingly worried about their health and nutrition. Wheat is the most extensively farmed cereal grain, and it meets the fundamental nutritional needs of virtually every food item. Wheat may be easily taken as a whole grain product and provides several health advantages due to the bioactive components found in its different layers (Fardet et al., 2010). Several studies have found that eating whole wheat has health advantages (Ounnas et al., 2014; Liu et al., 2018). The existence of anthocyanin-rich coloured wheat, on the other hand, has propelled whole wheat to the top of the list of health-promoting cereal grains (Liu et al., 2018; Sharma et al., 2020). Colored wheat with high anthocyanin concentration in the pericarp (purple wheat), the aleurone layer (blue wheat), or both (black wheat) may have a role in the prevention of oxidative stress-related illnesses (Garg et al., 2016; Sharma et al., 2018). However, coloured wheat germplasm is still in short supply, particularly those with high anthocyanin concentrations, acceptable processing quality, and high yield (Sharma et al., 2018). As a result, breeding coloured wheat with higher quality and yield characteristics has become a popular goal for increasing the value of processed wheat products. It has been discovered that coloured wheat lines are associated with a variety of unfavourable qualities that impact the yield trait, directly or indirectly. As a result, it is critical to have a thorough understanding of the pigmented wheat kernel genes and related features.

Many researchers studied and established that anthocyanin pigmentation in various organs of wheat is regulated by several regulatory genes like *R* (red grain), *Ra* (red auricle), and *Rc* (red coleoptile), *Pc* (purple culm), *Pg* (purple glume), *Plb* (purple leaf blade), *Pp* (purple pericarp), *Pan* (purple anther) (Khlestkina et al., 2008). However, the full molecular process underlying the coloration of purple pericarp, blue aleurone, and both layers of a wheat seed remains unknown. Wheat, being polyploid, anthocyanin regulation in its seeds is a complicated and high-priority process. Despite the fact that a few studies on purple wheat grain and blue wheat grains have sought to understand the process of colour generation, the needed depth is lacking (Liu et al., 2016; Jiang et al., 2018; Li et al., 2018; Jeewani et al., 2021). Global omics analysis is a significant resource for understanding the roles of key genes and regulatory mechanisms. In this work, we compared the transcript profiles of developing seeds of blue, purple, and black wheat with those of white wheat in order to acquire a deeper understanding of anthocyanin synthesis, regulation, and its impact on plant morphology.

## Results

### Agronomical trait evaluation

#### Seed parameters

Seeds collected at different days after anthesis (21, 24, 26, and 28 DAA) represented different seed development stages from milky ripe to dough ripe. At 21 DAA, seeds showed initiation of color accumulation, while at 28 DAA, seeds had maximum pigmentation (Figure 1A). The mature seed width was highest in purple wheat and followed the purple > white> blue >black trend. The seed length was most elevated in blue grain and followed the direction of blue>black>purple>white. The total kernel weight (TKW) was highest in purple wheat and followed the trend of purple>white>blue >black (Figure 1B). Grain hardness was highest in black wheat and followed the direction of black> purple> blue>white.

**Figure 1:**
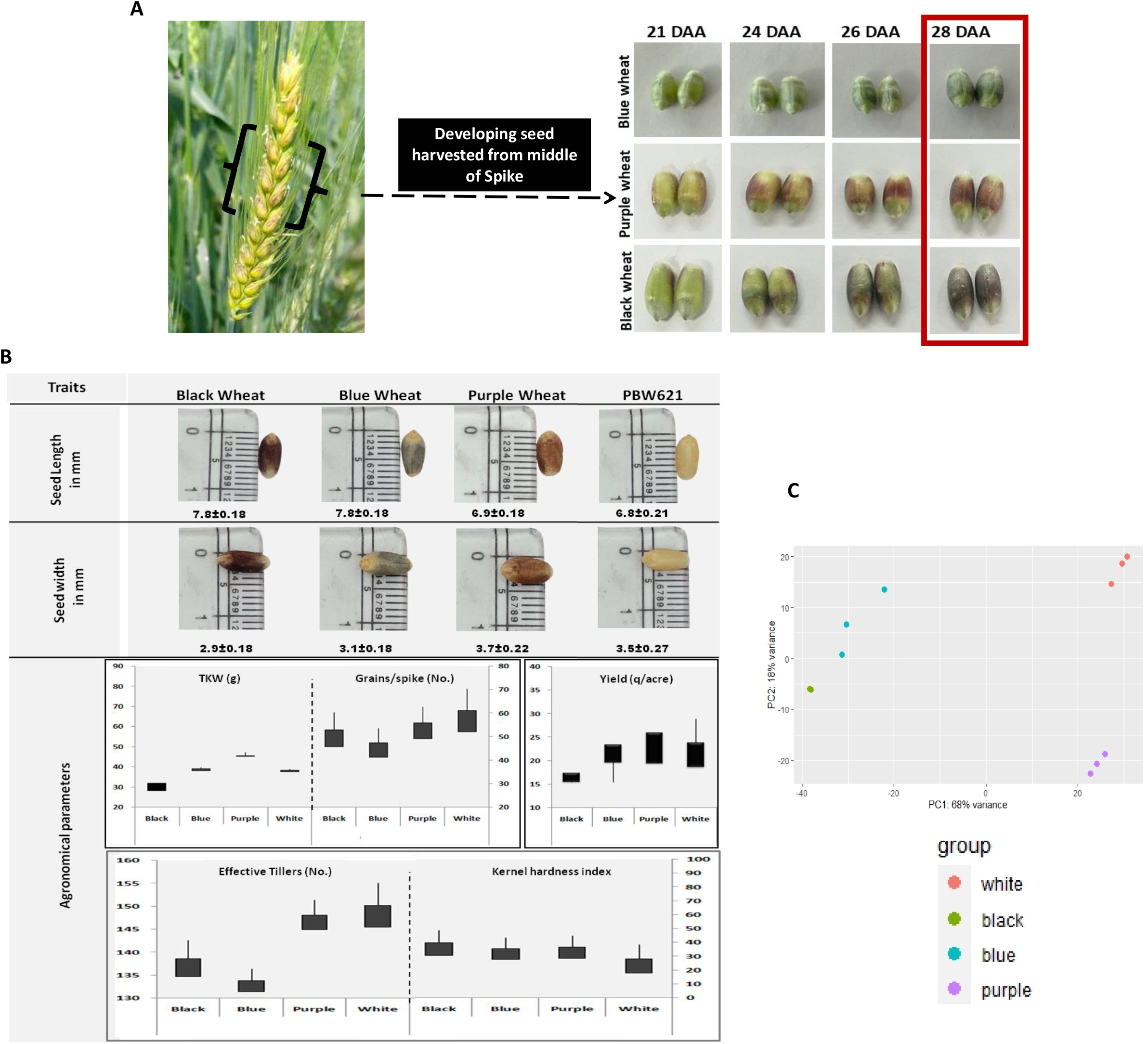
Pigmented seed morpho-physiological evaluation and transcriptomics profiling. **(A)** The pigmentation pattern of developing colored wheat seeds was examined at 21, 24, 26, and 28DAA. Spike at or after 28DAA i.e. 85 Zodak’s stage were collected, when the whole seed becomes pigmented. Seeds were dehulled from the middle of the spike and pooled for RNA-seq analysis. **(B)** Agro-morphological and related traits of matured colored and white wheat seed were examined. Wheat kernel size was considered by measuring seed length and seed width in millimeters. Yield influencing traits i.e. Thousand kernel weight (TKW) in grams, number of grains per spike, yield in quintal per acre, number of effective tillers per square meter, and kernel hardness index were measured. Data represents Mean + SD from 10 replicates. (C) Illustration of PCA plot to know the relationship between transcriptome data of all samples. PC1 (68% variance) and PC2 (18% variance) grouped black and blue wheat in close proximity, while purple and white wheat were grouped separately.

#### Crop Parameters

During the crop cultivation, the white wheat had the highest number of effective tillers, grains per spike, and yield that followed the trend of white > purple> black >blue (Figure 1B). The number of spikelets per spike was higher in white contrasted to others.

### Anthocyanin accumulation pattern

The amount and pattern of anthocyanin accumulation varied at different seed developmental stages and were light-dependent. The blue color accumulation started earlier than the purple color. Blue color initiated from the embryo and proceeded in the concentric rings along the length. In purple, it began from the ventral side towards the embryo and proceeded longitudinally (Figure 1A). The anthocyanin accumulation in the mature seeds was highest in black and followed the trend of black >blue >purple >white (Table S1). The genotype, environment, and the accumulating tissue (seed coat layer) influenced the anthocyanin accumulation pattern. Measurement of anthocyanins from two different locations and climate conditions showed that its higher accumulation occurred in the crop grown at high altitude and lower temperature (Table S1).

### Anthocyanin profiling by LC-MS/MS

The LC-MS/MS analysis was performed to determine the anthocyanins in the black, purple and blue wheat. The MS/MS spectra of anthocyanins were determined by using Electrospray Ionization (ESI) in positive mode. The findings revealed 26 different types of anthocyanins in black, 19 in blue, and 11 in purple wheat (Table 1). The confirmation of the anthocyanins was carried out by their molecular weight ion. The result showed that the mass to charge (m/z) ratio of different anthocyanins in the black, purple and blue wheat was in the range of 449-803. The anthocyanins compounds such as petunidin derivative (m/z 551 and 611), cyanidin derivative (m/z 565), peonidin di-glucoside (m/z 625), malvidin rutinoside (m/z 639), malvidin p-coumaroyl glucoside (m/z 639) are limited to only black wheat. Further, the black shared few anthocyanins with the blue wheat such as delphinidin glucoside (m/z 465), malvidin glucoside (m/z 493), petunidin acetyl pyranoside (m/z 521), peonidin derivative (m/z 579), cyanidin rutinoside (m/z 595), delphinidin rutinoside (m/z 611), cyanidin diglucoside (m/z 611), petunidin p-coumaroyl glucoside (m/z 625), and delphinidin diglucoside (m/z 627). In addition, ten anthocyanins were found common in black, purple, and blue wheat.

**Table 1:**
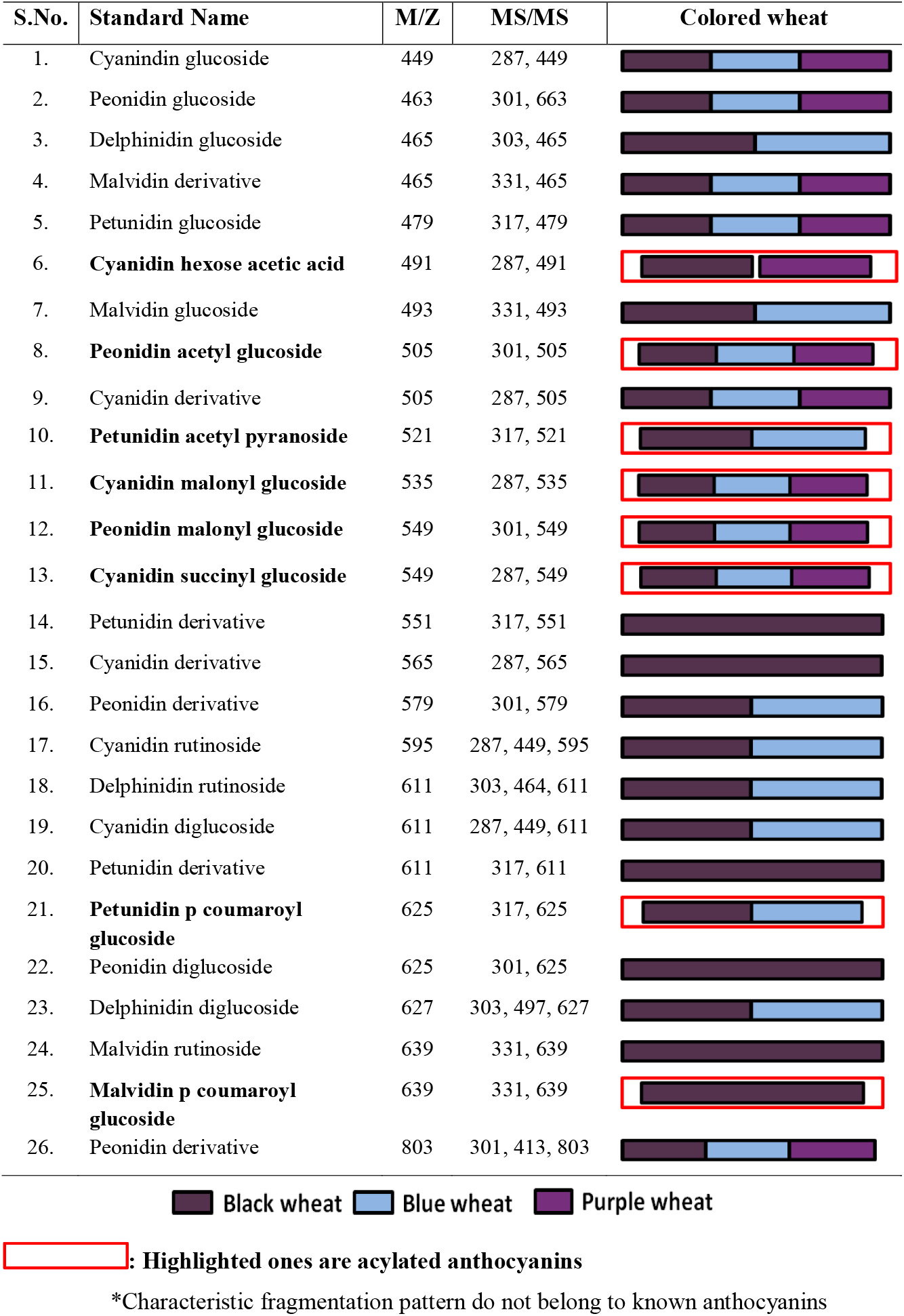
Composition of anthocyanin in colored wheat lines using LC-MS/MS.

As a part of complexing and co-pigmentation, anthocyanins can form esters with organic acids and other hydroxycinnamates. The few may contain acetyl, coumaroyl, and malonyl compounds (Stein-Chisholm et al., 2017). Our study reported all these acylations.

Furthermore, the presence of delphinidin derivatives was the leading cause of blue hue in blue aleurone, as supported by the previous report (Jeewani et al., 2021). As blue aleurone is present in both blue and black wheat, a similar remark had been noticed for both (Table 1). Although there were fewer anthocyanins in purple wheat, half (45%) were acylated, compared to 31% in black and 32% in blue wheat. Based on the finding, it was inferred that black grain has more anthocyanins of various types, in addition to a higher content hence maximizing its nutritional quality. On the other hand, Purple wheat has a high concentration of acylated anthocyanins, which have been extensively documented for their stability and bioavailability, and consequently various health advantages (Giust and Wrolstad, 2003; Zhu et al., 2018).

### Seed transcriptome profiling

Transcriptome profiling at the peak of anthocyanin accumulation (28 DAA or 85 Zadok scale) yielded 38.9 to 54.4 million raw reads per sample (Table S2). Filtered reads with Phred scores of 20 or above ranged from 37.6 to 52.2 million per sample (Table S2). Clean reads were uniquely mapped to the wheat genome, with more than 70% assigned to the genic region. The total number of expressed transcripts with TPM and RPKM >0 from all four samples was 85077.

PC 1 and PC 2 explained 68% and 18% variation in the PCA plot for diversity analysis, respectively. Wheat varieties were clustered into four distinct groups. The white wheat formed a distinct cluster separated away from the colored wheat. Among colored wheat, blue and black were positioned in proximity to each other, while purple lay distantly (Figure 1C). Black wheat had the least inter-replicate variance, followed by purple and white wheat, while blue wheat had the most inter-replicate variation.

Similarly, cluster analysis divided the four wheat lines into two primary clades. Black and blue were found in one clade, while purple and white were found in another (Figure S1). The PCA and cluster analysis both indicated a similar grouping structure.

### The transcriptional signature pattern of pigmented wheat seeds

The transcriptional signature associated with the pigmentation process was looked forward to analyzing differentially expressed genes (DEGs) in colored wheat in pairwise comparison with white wheat. The highest number of DEGs were observed in black wheat (1092), followed by blue (741) and then purple wheat (220) (Figure 2A). We hypothesized that black wheat had the combination of transcriptional signatures of blue and purple wheat. That hypothesis proved wrong when 464 DEGs were observed to be common between black and blue, 29 between purple and black, and 26 between purple and blue (Figure 2A). The observed unique expression pattern might be associated with other morphological, physiological, and rheological changes in the grain besides pigment production. Among all the three color wheat samples, 51 transcripts were observed to be common to white wheat. However, 548, 200 and 114 transcripts were unique to the black, blue, and purple wheat, respectively.

**Figure 2:**
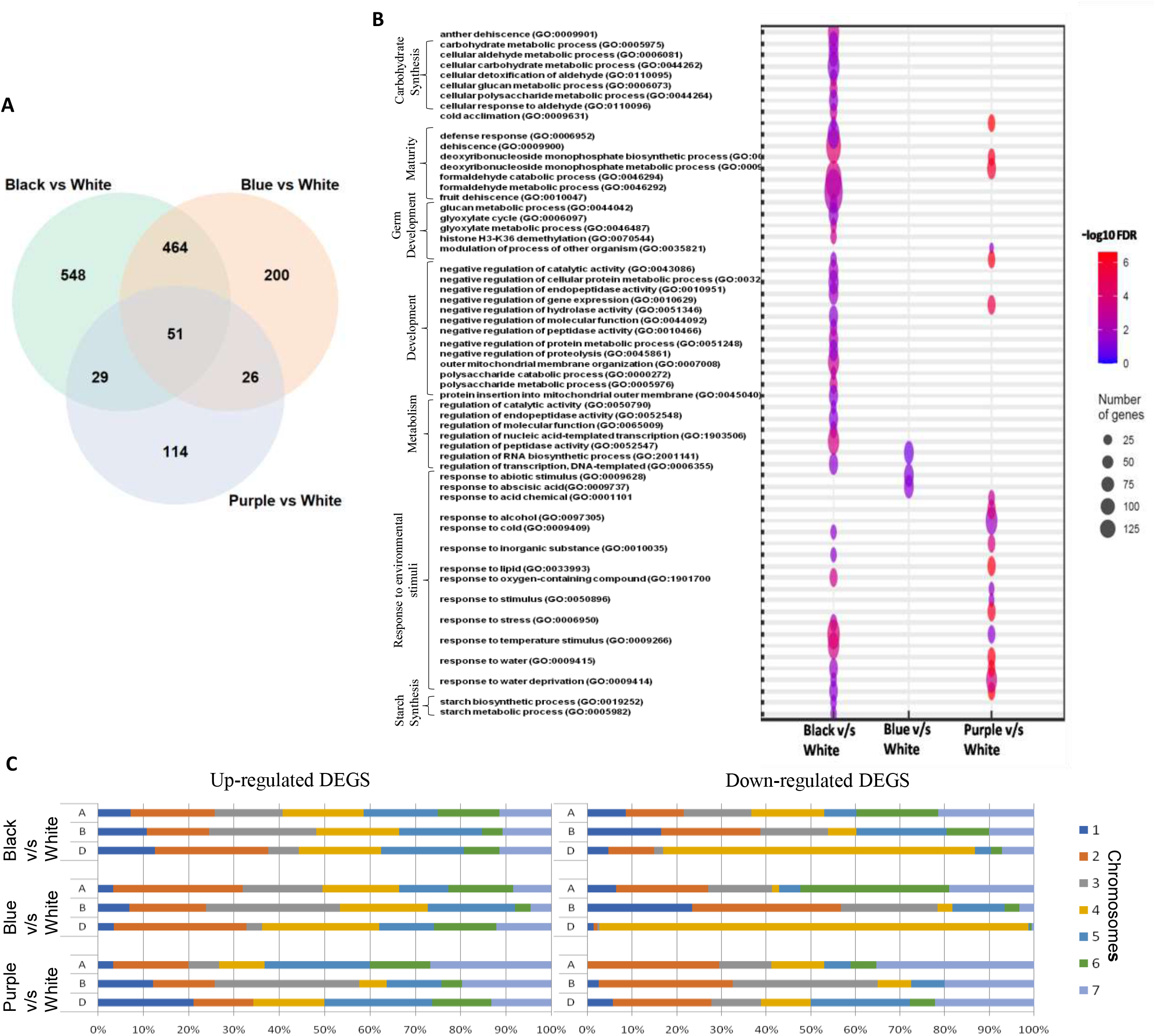
Transcriptional Signatures of three colored wheat seeds. **(A)** Venn Diagram showing the number of common and unique DEGs among three colored wheat samples compared to white wheat. Black wheat showed the highest number of DEGs and shared more DEGs with blue wheat in comparison to purple wheat. **(B)** Biological processes in GO enrichment analysis of DEGs. Only significantly enriched values with corrected *P* < 0.05 were included. The color and size of each point represented the -log10 (FDR) values and enrichment scores. A higher -log10 (FDR) value and enrichment score showed a greater enrichment. **(C)** Up and down-regulated DEGs across the A, B, and D genome of all chromosomes in studied colored wheat in comparison to white wheat. DEGs: Differentially Expressed Genes, GO: Gene Ontology, FDR: False Detection Rate

The chromosomal distribution pattern of DEGs showed a chief contribution of chromosome 4D in the black and blue wheat, while, in purple wheat, individual chromosome contribution was not so prominent (Figure S2). Further Gene Ontology (GO) term enrichment was performed to know DEGs’ biological relevance and functional significance (Figure 2B, Figure S3). It suggested that more biological processes were represented in black wheat followed by purple and, least, in blue. Enrichment of biological processes, including metabolic processes and their regulation, carbohydrate metabolism, germ cell development, processes relevant to maturity, depicted that the seeds were close to maturity at the selected developmental stage, and the endosperm was enriched with carbohydrates. Intriguingly, several biological processes linked to an environmental stimulus like a response to temperature, moisture, nutrients, etc., were prominent in black and purple wheat, with purple wheat showing a higher false detection rate than black. Whereas, processes relevant to starch synthesis were enriched only in black wheat. On looking at DEGs fold change value, we observed that black and blue wheat transcripts were more down-regulated than purple wheat (Table S3). Chromosome-wise expression value depicted that a maximum number of down-regulated transcripts belonged to the 4D in black and blue wheat (Figure 2C). However, up-regulated DEGs were almost equally distributed in all chromosomes of three colored wheat samples.

Cluster analysis carried out using top 2500 differentially expressed genes identified six clusters (Figure 3) that were further subjected to GO analysis. In clusters I and II, black and blue showed similar expression patterns, while the pattern of purple was like white wheat (Figure 3A and 3B). Genes in cluster I were enriched for processes broadly attributed to the cell wall and polysaccharide catabolic processes and response to abiotic and biotic stress (Figure 3A and Figure S4), which signified the attainment of seed maturity in black and blue wheat. Whereas cluster II was predominantly enriched for processes involved in core cell metabolic processes, development, and carbohydrate metabolism, especially starch biosynthesis (Figure 3B and Figure S4), which signified continued accumulation of carbohydrates in the endosperm of purple and white wheat. Clusters III and IV had an almost equivalent number of genes and similar expression patterns in all the colored grain relative to the white wheat (Figure 3C). Cluster III was enriched with processes involved in abiotic stress, and cluster IV was enriched with processes like secondary metabolites synthesis, including phenylpropanoid biosynthetic process, which directs towards anthocyanin accumulation in all the colored wheat lines (Figure 3C and Figure S4). Cluster V had a similar expression pattern in purple and black colored wheat relative to the white and blue wheat (Figure 3D). Cluster VI had a differential expression pattern in purple relative to the white, blue and black wheat (Figure 3D and Figure S4).

**Figure 3:**
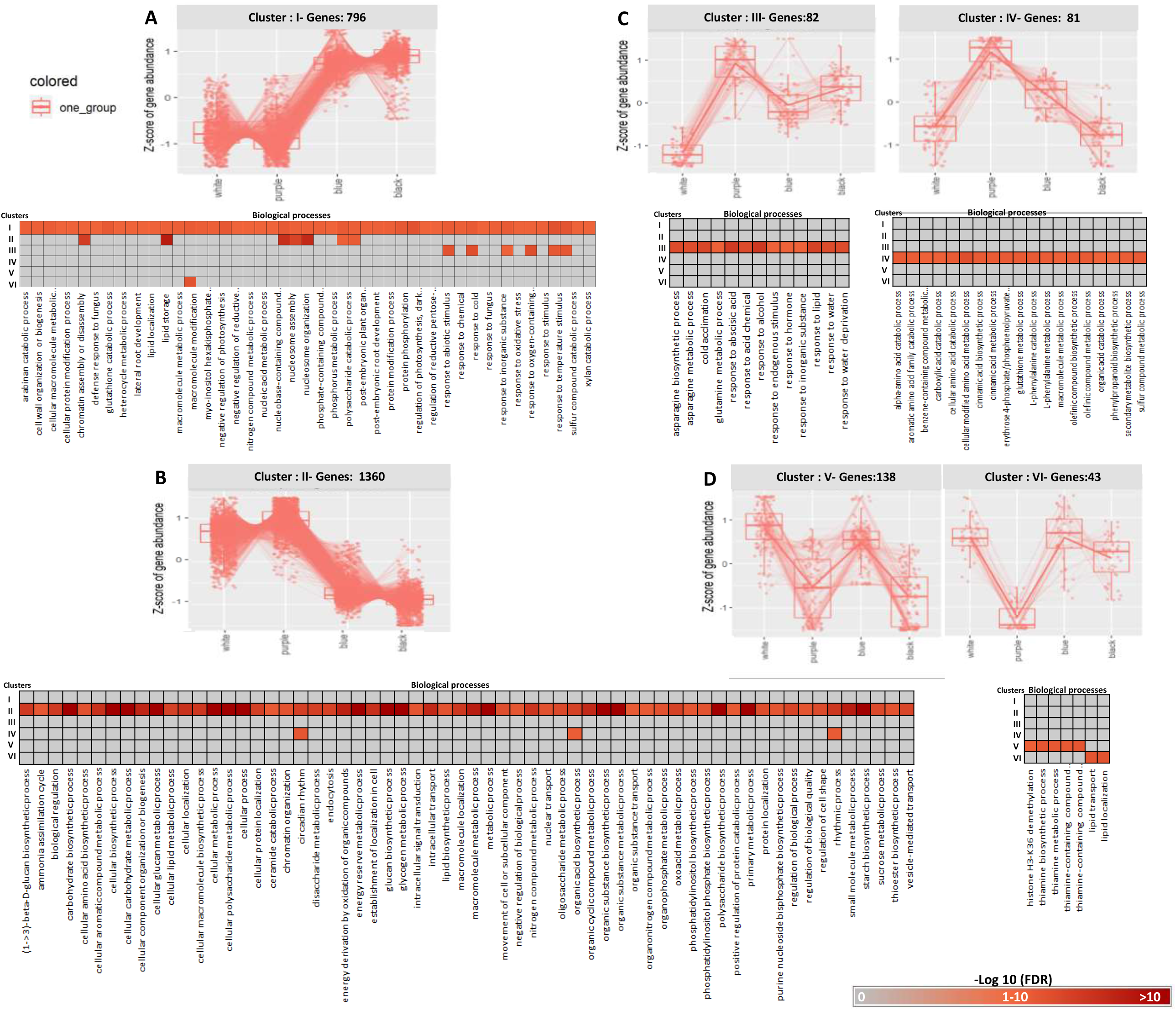
Co-expression clusters of DEGs inferred using hierarchical clustering. **(A)** to **(D)** Top 2500 DEGs with adjusted p-value were clustered into six groups with similar expression patterns across the different wheat samples and genes enriched for biological processes in respective clusters. **(A)** Cluster I represents 796 genes, broadly belonging to the cell wall and polysaccharide catabolic processes, and response to stress; where black and blue wheat showed a similar kind of expression pattern and, purple and white wheat alike. **(B)** Cluster II represented by the highest number of genes i.e.1360 and expression pattern wheat samples were exactly opposite to cluster I. It is enriched with the core cell metabolic processes, development, and carbohydrate metabolism processes. **(C)** Cluster III and IV correspond to an almost similar number of genes and expression patterns, where Cluster III represents processes relevant to stress and cluster IV is relevant to secondary metabolites. **(D)** Cluster V represented by 138 genes, where purple and black wheat showed a similar kind of expression pattern, and white and blue alike, enriched with thiamine and sulphur metabolic processes. Cluster VI is represented by the least number of genes i.e. 43, where purple wheat is expressed differentially as compared to others and enriched with lipid transport/localization processes.

Overall, based on transcriptional expression differences, we can conclude that black wheat showed more similarity with blue than purple. In both black and blue wheat, the carbohydrate metabolism and core metabolic processes were observed to be less expressed than purple and white wheat, which may contribute to the oblong seed shape in black and blue, and plump in purple and white wheat seed. Patterns common in black and purple wheat represented environment-related responses, including biotic and abiotic stresses indicating their role in plant adaptation.

### Expression of genes involved in anthocyanin biosynthesis and regulation

#### Regulatory Genes

For a comprehensive understanding of the expression pattern of anthocyanin regulatory and structural genes, DEGs were thoroughly studied for relative variation. Our data showed that three transcription factors (TFs) were activated in colored wheat samples compared to white wheat (Table S4). Among the identified TFs, two belonged to the *MYC* family and one to the *MYB* family. The first in the *MYC* family was from chromosome 4D and was highly expressed in the black, followed by blue wheat and not in purple wheat (TraesCS4D02G224600: 4D-*MYC;* anthocyanin regulatory R-S protein-like [*Aegilops tauschii* subsp. *strangulata*]) (Figure 4A; Table S4). The second in the *MYC* family was from chromosome 2A and was highly expressed in both blue and purple wheat (TraesCS2A02G409400: 2A-*MYC;* anthocyanin regulatory R-S protein-like isoform X1 [*Triticum dicoccoides*]) (Figure 4A; Table S7). The third in the *MYB* family was from chromosome 7D and was highly expressed in black, blue, and purple wheat (TraesCS7D02G166500: 7D-*MYB*; anthocyanin regulatory C1 protein [*Aegilops tauschii* subsp. *strangulata*]) (Figure 4A; Table S4). It should be noted that all the colored wheat varieties showed the expression of both kinds of transcription factors (*MYC* & *MYB*) i.e., at least one of the two differentially expressed *MYC*s. Interestingly, the expression of *MYB* was significantly higher in colored wheat (TPM 50-118), although low expression was detected in white wheat also (TPM 2.6) (Table S5). Out of two *MYC*s, 4D *MYC* showed TPM>500 in black and blue wheat, and 2A *MYC* showed TPM>1000 in purple and blue wheat. None of them were expressed in white wheat. TPM values showed these TFs are highly expressed and thus seem to regulate the anthocyanin biosynthesis. A similar expression pattern was also validated by qRT-PCR (Figure S5).

**Figure 4:**
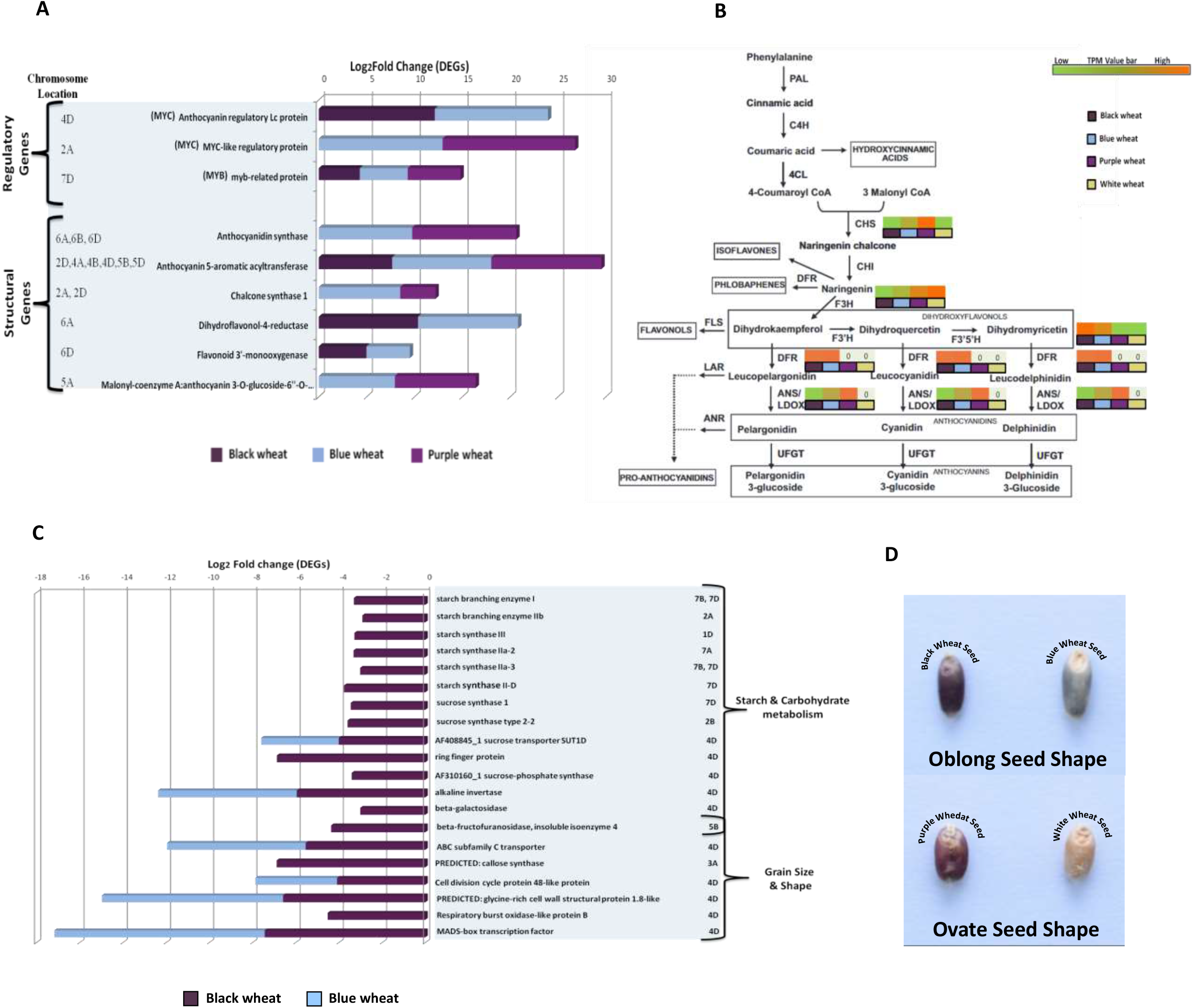
Expression pattern of Anthocyanin biosynthesis supportive genes and seed size trait. **(A)** Highly expressed genes of Anthocyanin regulatory and structural genes, where DEGs with > 3.5 log2 Fold value represented along with their chromosomal position. **(B)** TPM values of anthocyanin structural genes in phenylpropanoid pathway among three colored wheat lines and white wheat represented by a colored bar, where high TPM values were shown in red color and low TPM in green color. **(C)** Expression profile of seed size affecting DEGs with <-3 log2 Fold change along with their chromosomal position. **(D)** Colore wheat and white seeds image illustrating, black and blue wheat had oblong seed shape whereas purple and white wheat had obovate and plump.

Anthocyanin biosynthesis pathway is regulated by ternary complex (MBW) made from three families, i.e., *MYB*-bHLH-WD40, as shown by several researchers in different crops (Petroni and Tonelli, 2011; Patra et al., 2013). Previously Jiang et al. (2018) showed that in wheat *MYB* from chromosome 7D (TaPpm1; 7D) & *MYC* from chromosome 2A (TaPpb1; 2A) co-regulate anthocyanin synthesis in the purple pericarp through the MBW complex. At the same time, Liu et al. (2016) has documented *MYB* from chromosome 7B (TaPpm1; 7BL) & *MYC* from chromosome 2A (TaPpb1; 2AL) for purple wheat. In the current study, putative *MYB* related protein (TraesCS7D02G166500: 7D-*MYB*) was identical to TaPpm1 of Jiang et al. (2018) and was confirmed through multiple alignments and conserved motif analysis by MAST (Figure S6 & S7). Whereas, two *MYC* TFs viz., TraesCS2A02G409400: 2A-*MYC* and TraesCS4D02G224600: 4D-*MYC* had identical bHLH motif to TaPpb1 (Figure S6 & S6). TaPpb1 for purple pericarp was observed to be more identical to 2A-*MYC* in multiple alignments (Figure S6). Thus, our study aligns with Jiang et al. (2018) observations.

Previously Li et al. (2017) showed that blue color requires *ThMYC4E* (4E (4A/4B/4D) to regulate anthocyanin synthesis in the aleurone of wheat. 4D-MYC observed in black and blue in our case represents this TF. Our observations indicated that 4D-MYC and 7D-MYB co-regulated the blue aleurone coloration. Out of 743 TaWD40 proteins reported in wheat (Hu et al., 2018), 394 observed in the current study exhibited almost similar expression patterns in all colored and white wheat seed transcripts indicating that WD40 is expressed in wheat seed irrespective of its color (Table S6). Thus, purple coloration is controlled by 2A-MYC, 7D-MYB, and WD40, while blue aleuron by 4D-MYC, 2A-MYC, 7D-MYB, and WD40.

#### Structural Genes

Exploration of structural genes indicated that the TPM values of all of them were higher in colored wheat than white. Those with high fold differentially expressed were thoroughly studied. It indicated that *F3’H* and *DFR* were highly expressed in black and blue wheat and *ANS* in blue and purple wheat. *ANS* isoforms were also identified. Two isoforms were observed in blue and 3 in purple. *CHS* was highly expressed in both blue and purple wheat (Figure 4A & B). A high-fold expression of genes involved in the acylation of anthocyanins was also observed. *O-malonyltransferase* was highly expressed in blue and purple wheat, which transfers the malonyl group to anthocyanins to produce anthocyanidin 3-O-6’’-O-malonyl glucosides (Figure 4A). Another important acylating gene, i.e., *5-aromatic acyltransferase, was* highly up-regulated in purple followed by blue and black wheat, which transfers hydroxycinnamic moieties to the glucosyl groups of anthocyanin (Figure 4A). All highly expressed structural genes and their isoforms localized to 2A, 2D, 4A, 4B, 4D, 5A, 5B, 5D, 6A, 6B, 6D chromosomes. It indicated that structural genes were not localized to a single place but instead distributed to different chromosomes (Figure 4A).

Besides anthocyanin biosynthesis, other genes like *High molecular mass early light-inducible protein HV58, O-methyltransferase ZRP4, Os10g0395400* (involved in anthocyanin transportation in black rice caryopsis), and *MATE efflux family* indirectly known to enhance the anthocyanin accumulation were observed to be differentially expressed in purple, blue and black wheat, respectively (Table S7).

### Expression of genes involved in seed morphology, physiology, and rheology

Our dataset facilitated the recognition of genes with characteristic expression in pigmented wheat seeds compared to un-pigmented wheat (Table S7 & Figure S8). To explore the patterns and expression of genes associated with seed development, DEGs were further explored. The soft dough stage (28 DAA) of the wheat kernel is the last step of kernel development, in which cell division and extension go on for starch deposition, vital for kernel development and size (Baroja-Fernández et al., 2003; Daba et al., 2020). However, in our dataset, we observed that most of the significant genes viz., *starch branching (sb) I, sbIIb, starch synthase (ss)III, ssIIa-2, ssIIa-3, ssII-D* encoding enzymes of starch synthesis were highly down-regulated in black wheat (Figure 4C). Besides that, the *ring finger protein* gene, which plays a role in the starch accumulation and wheat seed size (Parveen et al., 2021; Elhadi et al., 2021), was down-regulated only in blue wheat (Figure 4C). Sucrose breakdown is very important for starch biosynthesis, as sucrose is the major carbohydrate source for seed development (Baroja-Fernández et al., 2003; Daba et al., 2020). Notably, *sucrose transporter SUT1D; sucrose-phosphate synthase; alkaline invertase; beta-fructofuranosidase; insoluble isoenzyme 4* was highly down-regulated either or both in black and blue wheat, respectively (Figure 4C & Table S7). Moreover, *ABC subfamily C transporter*; *callose synthase*; *cell division cycle protein 48-like protein, glycine-rich cell wall structural protein 1*.*8-like*; *respiratory burst oxidase-like protein B*; *MADS-box TF* genes which were reported to be directly or indirectly regulating the grain size by endosperm development, cell size expansion and growth (Table S7) (Walter et al., 2015; Cheng et al., 2017; Ma et al., 2017; Daba et al., 2020) were also highly down-regulated either or both in black and blue wheat, respectively (Figure 4C). No significant changes were seen in purple wheat compared to the white wheat kernel (Table S7). Captivatingly, most of the smaller seed size-related genes were localized in 4D chromosomes of black and blue wheat (Figure 4C). Morphological observations also indicated that blue and black wheat had similar oblong seed shape as compared to obovate and plump seeds of white and purple wheat (visual confirmation in Figure 4D).

The storage protein genes, including LMW-glutenin subunits, α/β/γ/ω-gliadins, *avenin-like protein*, and *puroindoline-A*, were more expressed (log2FC<5) in colored wheat seeds than white wheat (Table S7). That may be attributed to the final grain protein content.

### Chromosome specific characterization using SSR markers

Advanced colored wheat lines used in this study were developed by backcrossing the exotic winter wheat lines, i.e., BW (donor parent) with Indian high yielding released spring wheat cultivar PBW621 (recipient parent). These lines were screened by genome-wide SSR markers to find out the recipient parent restoration. We got polymorphic amplification profiles of 101 markers in wheat lines among 166 SSR markers screened (Table S8). Marker data suggested 66.66%, 67.5%, and 77% background recovery in black, blue, and purple wheat, respectively (Figure S9). Better background recovery was observed in chromosomes 4 (except 4D), 5, and 6 (Figure S9). As the donor parent was a 4E substitution line, additional 36 consensus SSR markers were employed to assess the involvement of *Thinopyrum* chromosome 4E in color wheat (Table S9 & S10). The 4D chromosome in black wheat was found to be about 85 % more similar to the donor, followed by 73 % in blue wheat and the least in purple wheat (10 %) (Figure 5A).

**Figure 5:**
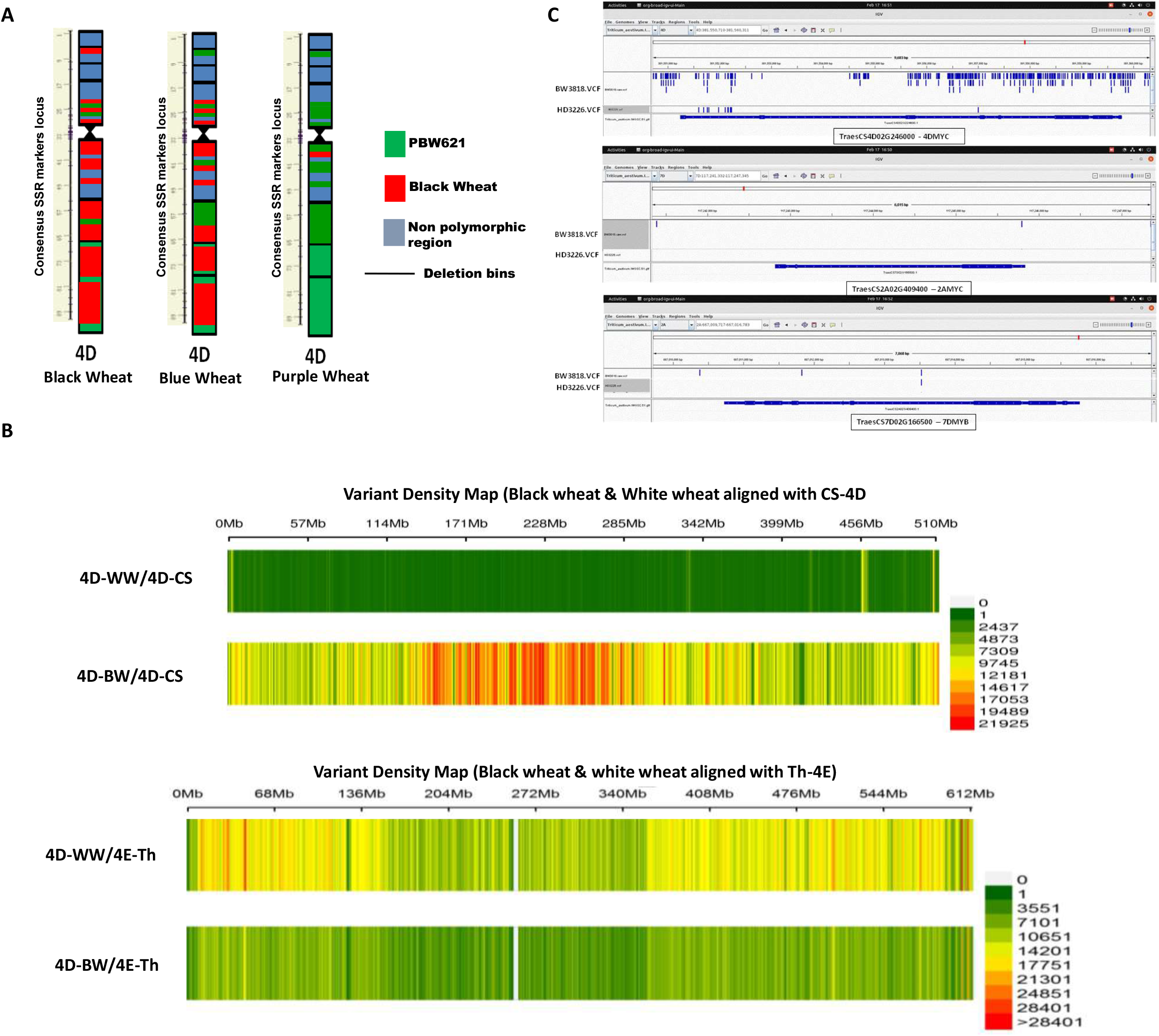
Genomic overview of advanced colored wheat lines. **(A)** Schematic representation of consensus SSR marker based map of 4D chromosome of black, blue and purple wheat; color represented the parent similarity, PBW621 recipient parent (green), BW donor parent (red), and blue as non-polymorphic regions. **(B)** Chromosome-wise distribution of genic variants in 4D chromosome of black (4D-BW) and white wheat (4D-WW) in reference to chinese spring 4D (4D-CS) chromosome and *Thinopyrum elongatum* 4E (4E-Th) chromosome. **(C)** Comparative visualization of 4D-MYC, 2A-MYC and 7D-MYB using the IGV browser in black and white wheat. VCF files of black and white wheat uploaded along with white wheat reference genome (IWGC).

### Genomic variation in black wheat

The incidence of variations in genes and genomic regions provide important information for understanding the genomic makeup of the concerned organism. Thus, we compared the whole genome sequences of black and white wheat with the reference genome of Chinese Spring (CS) wheat to determine the distribution pattern of all genomic variations (SNPs, Indels, and complex nucleotide variants). It was seen that the 4D chromosome of black wheat showed very high variation compared to CS, while the 4D chromosome of white wheat showed the least variation (Figure S10). At the same time, the rest of the chromosomes had a similar variation pattern in black and white wheat. Further, to pinpoint recombination regions in this chromosome only, we examined the genomic variation distribution patterns of black and white wheat 4D chromosomes compared to CS-4D and Th.elongatum-4E chromosomes. It suggested a close link between the black wheat 4D chromosome and *Th*.*elongatum*-4E (Figure 5B).

Further, within the chromosome, proximal regions of both the arms of the black wheat 4D chromosome exhibited minimal variation compared to *Th. elongatum*-4E (Figure 5B), while the distal region (towards the centromere) displayed variations in small areas of a few Mbs. Interestingly, the proximal part of the 4D chromosome of white wheat also showed comparatively more minor variation than *Th. elongatum*-4E, ruling out translocation involving significant chromosome parts. It seems that the substitution chromosome (4E{4DL}) obtained from the donor parent is consecutively changing with very small recombinations in every generation.

Moreover, on visualization of some selected genes like putative TFs from transcriptional studies, in the IGV browser revealed that black wheat-4D derived gene has a higher number of variants in comparison to white wheat-4D (Figure 5C).

## Discussion

The transcriptional signatures and molecular circuitries in this study revealed that colored wheat lines were not only different in terms of grain color but for several other traits. This comprehensive transcriptomics study of different colored wheat seeds (black, blue, and purple) in comparison to white wheat at the dough developmental stage suggested that the anthocyanin regulatory and structural genes expression program work almost similarly and independent of tissue localization, whether the pigmentation is in pericarp or the aleurone layers of wheat seed. We found at least two anthocyanin regulatory transcription factors, i.e., MYB and MYC (bHLH), are required for the anthocyanin biosynthesis in pericarp and/or aleurone layers. However, the transcriptional signature pattern was observed to vary with the pigment localization in the seed, which might be linked with the allelic variation and chromosome localization of the concerned gene. The small seed size of black and blue wheat was because of the linkage drag associated with the 4E chromosome of *Th*.*elongatum* that replaced the 4D chromosome of wheat. The transcriptional evidence also highlighted the alliance of pericarp-associated grain pigmentation trait with response to abiotic stresses, including temperature, drought, and UV irradiation.

Previous research has found that MYB and MYC TFs are required for purple pericarp color in wheat (Koes et al., 2005; Jiang et al., 2018, Li et al., 2018). Whereas a single TF, ThMYC4E, has been described in blue wheat (Li et al., 2017). The most notable point in our transcriptome data set was that both MYB and MYC TFs are necessary to control the expression of structural genes involved in anthocyanin biosynthesis via a ternary complex. 2A-MYC and 7D-MYB controlled purple wheat pericarp pigmentation, whereas blue wheat aleuron pigmentation was regulated by three TF, including two MYCs (4D-MYC and 2A-MYC) and one MYB (7D-MYB). WD 40, on the other hand, was discovered to express non-differentially, i.e., in pigmented as well in white wheat. In contrast, an intriguing observation was made in black wheat, where two TFs, 4D-MYC, and 7D-MYB, controlled anthocyanin production. Our observations have been supported by several other studies on different crops (Procissi et al., 1997; Petroni and Tonelli, 2011). Both TFs have been shown to have a significant allelic diversity, with many homologous genes that regulate anthocyanin synthesis in a tissue-specific manner (Procissi et al., 1997; Petroni and Tonelli, 2011; Zong et al., 2017 and Jiang et al., 2018). Even in previous genetics research articles, the expression of the purple pericarp trait was reported to be regulated by two complementary factors coding for TFs (Gordeeva et al., 2015), which are present in separate genomes, namely A and B or A and D (Tereshchenko et al., 2012). This observation undoubtedly seems true in purple wheat (A and D genomes). However, black wheat color was controlled by two TFs from the D-genome and blue wheat color by three TFs from the A and D-genomes. It suggests that a single MYC, in conjunction with an MYB, can trigger anthocyanin production in wheat pericarp and aleurone. Two *MYCs* found in blue wheat may be different isoforms that complement each other. Since MYB and WD40 have a broad expression, MYC seems seed-specific (Jiang et al., 2018). It shows that MYC introgression is vital for grain color development in wheat and that fundamental machinery (MYB and WD40) is required to form a ternary complex that governs anthocyanin production in wheat.

Previous research has found that a temperature-responsive region in the MYB promoter regulates anthocyanin formation in apple, cotton, and blood orange (Espley et al., 2009; Gao et al., 2013; Huang et al., 2019). Besides that, *MYB* has also been reported to limit anthocyanin accumulation (Procissi et al., 1997) because it is further controlled by light intensity or UV irradiation (Procissi et al., 1997; Shin et al., 2013; Jiang et al., 2018). This scrutiny also proved true in our studies, as colored wheat lines purple, black and blue wheat showed almost 2.5, 1.7, and 1.5 fold higher anthocyanin accumulation, respectively, at high altitude areas with cold climate and high UV irradiation (Table S4). Another probability might exist that some other light regulating TFs in combination with *MYB* and *MYC* enhances the pigmentation in the wheat seed. Which still needs to be explored in wheat.

In addition to regulatory genes, we discovered heterogeneity in the expression pattern of anthocyanin structural genes in black, blue, and purple wheat seeds (Figure 4B) (Shoeva et al., 2014; Tereshchenko et al., 2013; Li et al., 2018). The expression of all structural genes was not uniform across developing-colored wheat seeds but instead followed a specific pattern with significant expression in either purple and blue or blue and black. *CHS* and *ANS* were highly expressed in purple and blue wheat, whereas *F3’H* and *DFR* in blue and black wheat. A similar expression has been reported by Li et al., (2018). Expression of these genes may vary with seed development, e.g. *CHS* might have accumulated earlier in black wheat. *CHS* was reported to be expressed 5 days prior to grain pigmentation and end prematurely (Trojan et al., 2014; Li et al., 2018), while *F3’H* expression decreased with the development stages (Khlestkina et al., 2008; Li et al., 2018). Though a common transcriptional regulatory program regulates all structural genes, they are still expressed differentially, depicting their frequency of expression in response to developmental stage and environmental changes (Koes et al., 2005 and Li et al., 2018). We also observed differential expression of anthocyanin modifiers (e.g. acyltransferase) in colored wheat (Li et al., 2018). Previously, Garg et al., and Li et al., [2016,and 2018] indicated that purple wheat had higher levels of acylated anthocyanins, which was confirmed in our samples since purple wheat transcriptome and LC-MS/MS data revealed higher levels of acylating enzymes and acylated anthocyanins, respectively.

Pigmentation in seed not only changes the appearance and nutritional quality of seed but protects it from biotic and abiotic stresses like insect attack, microbial pathogens, adverse temperatures, and UV-irradiations (Wan et al., 2020). Likewise, our gene enrichment results depicted 13 biological processes mainly associated with environmental simulative responses that were differentially altered in colored wheat seeds. Interestingly, it has been observed that anthocyanin presence in the outer layer, i.e. pericarp of purple and black wheat, influences the abiotic and biotic responses. Here, we can propose that these traits are more linked to *MYB* like TFs, which were stimulated by UV-irradiations (Wang et al., 2018) and cold stress (Procissi et al., 1997; Wan et al., 2020). Further, it has been reported by several researchers that pigmentation in wheat seeds was negatively correlated with grain yield, which was due to linkage drag associated with alien chromosome in color wheat (Martinek et al., 2014; Garg et al., 2016). Blue aleurone trait is contributed by the 4E that substituted 4D wheat chromosome. We observed that the substituted 4D part negatively regulated the carbohydrate metabolism and thus suppressed the synthesis of the carbohydrate and starch. It’s a well-known fact that enhanced starch synthesis in wheat and maize grain positively correlated with grain yield (Irshad et al., 2019; Daba et al., 2020). Gene enrichment also depicted other genes and factors that directly or indirectly determine the grain size affected by the 4E part.

Based upon the reduced expressions of the genes belonging to the 4D, SSR molecular marker mapping, variant calling using genome sequencing and IGV browser gene structure visualization, it was inferred that the advanced black and blue wheat lines used in this study were substitution line (4E{4D}), with some recombination regions in the distal part of 4E substituted 4D chromosome. That could be why these black and blue wheat lines perform better than usual full chromosome substitution lines in terms of agronomic performance, including yield and plant morphology. But to surpass the high-yielding white wheat cultivars, a substantial number of recombinations are still required. This might be reasonably possible in the coming years as the distal ends of the chromosomes have markedly higher recombination rates (Ramírez-González et al., 2018). On the other hand, purple wheat performs extremely well agronomically and is equivalent to high-yielding white wheat cultivars as it is only a mutant. These interpreted facts and observations will be valuable for future research into pigmented wheat and other grains system biology and breeding activities.

## Material and Methods

### Morphological observation of Seeds

In a completely randomised design, colour wheat advanced lines (BC3F8; with high anthocyanin content) [(Sharma et al., 2018) (pedigree given below) and common amber wheat (cv. PBW621) were cultivated in the experimental field at National Agri-food Biotechnology Institute (NABI), Mohali (30°44′10” N latitude at an elevation of 351 m above sea level) and Keylong, Lahaul Spiti. Seed length, seed breadth, total kernel weight, grains/spike, effective number of tillers, grain yield (q/acre), and kernel hardness index at maturity were studied and recorded for two consecutive seasons (2017-2018 & 2018-2019). Observations for morphological parameters were taken from ten plants chosen at random. Pedigree of advanced color wheat lines selected for RNA isolation

| Sample Name | Pedigree |
| --- | --- |
| White Wheat | PBW 621 (India-Punjab released variety) |
| Purple Wheat | BW/3*621 |
| Blue Wheat | BW/3*621 |
| Black Wheat | BW/3*621 |

BW is black wheat exotic black color donor line

### Anthocyanin extraction and estimation

Method for anthocyanin extraction and its estimation followed as shown by Kumari et al., (2020). Briefly, extraction was proceeded using acidified methanol (85:15-Methanol and 1N HCL) and stored at −20 °C after filtration with syringe filters of 0.45 µm pore size.

Anthocyanin content was measured by using spectrophotometer absorbance at 535 nm and total anthocyanin content was calculated and expressed as mg/Kg by using formula as described by Kumari et al., (2020).

### Anthocyanin composition measured by LC-MS/MS analysis

A Waters UPLC (Waters Corp., Milford, MA, USA) with an AB Triple TOF 5600 plus System was used for the MS analysis (AB SCIEX, Framingham, MA, USA). The optimum MS conditions were as follows: the scan range was set to m/z 100–2000, the source temperature was 550°C in positive ionisation mode, the pressure of Gas 1 (N2) and Gas 2 (N2) was 50 psi, and the curtain gas was set at 30 psi. The collision energy for MS/MS was 35 volts, the collision energy spread was 15 volts, and the declustering potential was 50 volts. The eluents A and B were 5% (v/v) formic acid and 100% acetonitrile, respectively. The injection volume and flow rate was set at 7 μl and 0.3ml/min. Chromatographic separations were done on a gradient program as follow: 0 – 5 min 95% A, 5 – 5.1 min 85% A, 5.1 – 6.1 min 84.5% A, 6.1 – 7min 0% A, 7.1 – 8 min 97% A and 8 – 15 min 5% A. Data processing was performed using Analyst TF software (version 1.5.1) and anthocyanins matching was carried out using Peak view software (version 1.2).

### Background screening of advanced colored wheat lines derived from a exotic substitution 4D (4E) line

CTAB method was used to isolate genomic DNA from fresh leaves. Quality of DNA was checked using 0.8% agarose gel and Qubit 3.0 fluorometer (Thermo-Fisher Scientific-Invitrogen). Black, blue, purple and white wheat DNA was used for background screening using a set of deletion bin-based SSR primers. While black and white wheat DNA was used for genome sequencing for estimating 4E and 4D translocation. For background recovery estimation, 166 SSR markers representing all wheat chromosomes were selected from the Sourdille map (Table S8 and S9) (Sourdille et al., 2004). In addition to above SSR markers, 4D specific, 36 SSR additional markers were used (Table S9 & S10).

### Sample collection for RNA Seq

For studying the pigmentation pattern in wheat seed, developing color wheat seeds in 28DAA (Day after Anthesis) or Zadok scale of 85 (soft dough, http://www.usask.ca/agriculture/plantsci/winter_cereals/winter-wheat-production-manual/chapter-10.php) were collected at 30°C day/17°C to 34°C day/18°C (day/night) temperature, when almost whole seed was pigmented (Figure 1A) from NABI fields. In brief, on the day of anthesis, the primary tiller was tagged. Samples were collected at four different DAA i.e. 21 DAA, 24 DAA, 26 DAA, and 28DAA. Complete pigmentation was seen on or after 28DAA of the wheat seed coat, thus this stage was selected for transcriptomics studies. 10 Spikes from three different plants were pooled together for one biological replicate in each sample and stored at -80° C for further studies.

### RNA isolation

Seeds were dehulled and immediately, RNA was extracted, using Sigma Spectrum plant’s total RNA kit. Three biological replicates from each sample were subjected to RNA-seq analysis.

### Sequencing

RNA and DNA samples passed by quality check were subjected to outsourcing library preparation and Illumina sequencing from Nucleome Informatics Pvt Ltd, Hyderabad with methodology, as shown in Figure S11 and S12. Four libraries were pooled per sequencing lane for Illumina HiSeq 2000 sequencing.

### RNA Seq

#### Mapping to Ref Genome and DEGs analysis

Final RNA-seq reads obtained after pre-processing of adapter sequences were processed for a quality check by using Fast QC. version0.11.5, http://www.bioinformatics.babraham.ac.uk/projects/fastqc/). They were filtered for low-quality reads (Phred Score ≤ 20) and short reads (length ≤ 20 bps) were eliminated. These reads were assembled and then mapped to the entire IWGSC RefSeq v1.1 reference genome. Clean RNA reads were mapped using STAR v2.5.3a (Dobin et al., 2013) with default parameters to generate gene-level counts using annotated reference genome GTF file. The normalized expression values of genes in all samples were estimated by salmon and summed by tximport and further used input in differential expression analysis.

Here, we generated results of differentially expressed (DE) genes by applying the likelihood ratio test on full formula ∼ phenotype and reduced formula as intercept only. Using an adjusted p-value cutoff of 0.001, obtained genes were ordered according to p-values and took the top 2500 for clustering to find genes with similar expression patterns.

Meanwhile, it has been observed that the black wheat 3rd replicate was grouping with blue samples in PCA plots, dendrograms, and LRT results. The results of Likelihood ratio test (LRT) and SERE (Schulze et al., 2012) were performed by using all replicates or by removing blue3, black3 or both. Thus, black wheat 3rd replicate was removed for separation of blue and black on dendrogram and PCA plots.

Results generated by differential expression analysis using Wald’s test followed by hypothesis testing to get DE genes (DE = log2fC > |2|; adj p-value< 0.05). White wheat was taken as control here for comparison.

For Statistical analysis, the R package and Microsoft Excel were used to analyze the data.

### Functional annotation, KEGG, and GO identification and their classification

DEGs were mapped to KEGG orthology ids using BLAST Koala. Because BLAST Koala did not annotate the whole DEGs of wheat, they were remapped using the NR database. Similar 20k DE genes (adjusted p-value< 0.001) identified by the LRT test were further mapped to rice Ids on KEGG using KOBAS. This mapping was done because KEGG does not support wheat. Only the first isoform of each DEG gene was kept. If multiple wheat Ids map to a single rice KEGG ids, then the values are averaged across the multiple wheat Ids. 20,000 wheat DE genes mapped to around 9000 rice KEGG Ids. R code was used to generate a matrix of R log values of differentially expressed genes along with their rice KEGG identifier.

### DEGs and further transcripts in silico functional analysis

Other than DEGs, transcript levels were further calculated using the TPM method (Transcripts per kilobase Million) and genes sorted with P≤0.01, TPM≥2. The anthocyanin-related genes selected from the DEGs were NCBI blasted with NR database to annotate maximum possible IDs. Putative TFs were further analysed using multiple alignments by Clustalvis and conserved motifs were identified using the Meme tool (http://meme-suite.org/tools/meme) with default settings except for the motif dimensions which were kept at, minimum width 5 and maximum width 50 amino acid (Kumar et al., 2020).

### Validation by qRT- PCR

Expression of Anthocyanin regulatory genes were validated by qRT-PCR. Relative fold expression changes and standard error were calculated using the 2-ΔΔCt method (Schmittgen et al., 2008) using three biological and technical replicates. Each cDNA sample was normalized with three endogenous reference genes i.e. GAPDH, ARF, and tubulin.

### DNA Seq

#### Data pre-processing and mapping to reference genome

Obtained raw sequencing files were pre-processed to remove adaptor sequences, low quality reads and below 90% accuracy rate using FastQC tool kit (Wingett and Andrews, 2018) https://github.com/OpenGene/fastp. High quality reads from two replicates were mapped to the reference genome (Ensembl *Triticum aestivum* release 52) using BWA-MEM tool (Durbin and Li, 2010) followed by samtools (version 1.10) for indexing of bam generated files. Samtools mpileup algorithm was used to combine mapped replicates in one file and visualize the file using VCF tool (Danecek et al., 2021).

### Variant calling and IGV visualization

Further, VCF tools position filtering options and Integrative Genomic View (IGV) (Robinson et al., 2017) were used to extract and visualise genomic variations (SNPs, InDels, etc.). Extracted variants were represented using the CMplot tool in R (Yin et al., 2021). Variation among chosen genes was also observed using the IGV window. Initially, the whole black and white wheat genomes were aligned and mapped with the reference genome Chinese Spring(CS-IWGC), however after obtaining the greatest number of variations in 4D chromosome, one chromosome (only 4D) genomic sequence was retrieved from sorted bam files of black and white wheat. They were remapped to the 4D and 4E chromosomes of the reference genome CS and *Th. elongatum* (https://www.ncbi.nlm.nih.gov/search/all/?term=Thinopyrum%20elongatum), respectively.

## Author contributions

SS and MG planned and designed the study. SS wrote the manuscript. SS, AK, and DS performed the experiments and analysed the data. AK, PK, SK, and BS helped in data sorting and manuscript editing. MG supervised the experiments and edited the manuscript. All authors contributed to the article and approved the submitted version.

## Acknowledgments

The authors are thankful to Dr. Humira Sonah for providing valuable data analysis advice. Authors are also thankful to High-Performance Computing (HPC) Cluster and Computational Biology Lab Facility provided by NABI-CDAC collaboration and DeLcO library facility. The authors declare that the research was conducted in the absence of any commercial or financial relationships that could be construed as a potential conflict of interest.

