## Supplementary figures for "Transcriptional signature pattern in black, blue and purple wheat and impact on seed pigmentation and other associated features: Comparative transcriptomics, genomics and metabolite profiling"

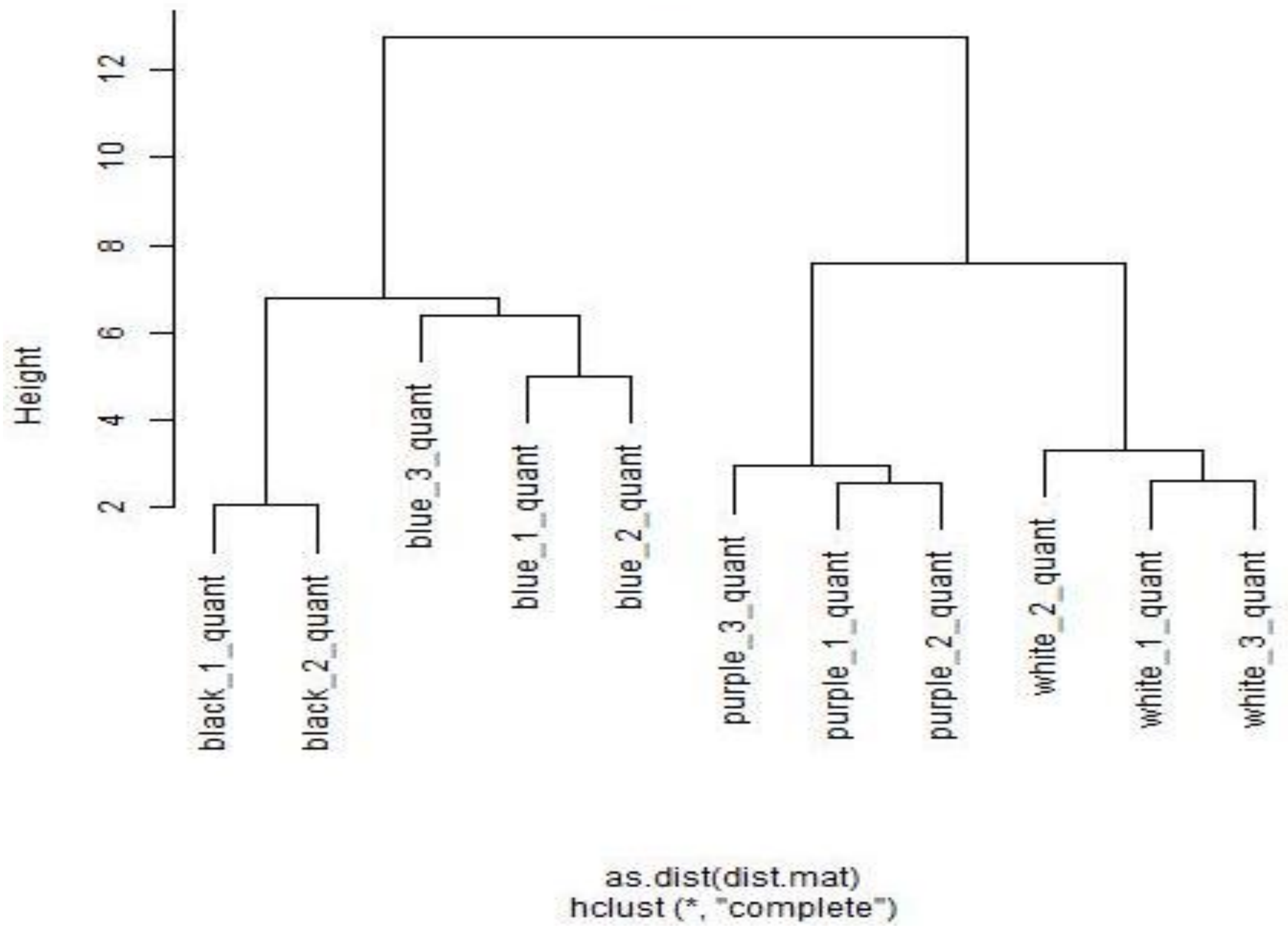


**Supplemental Figure 1: Cluster Dendrogram grouping replicates of samples into one cluster.**

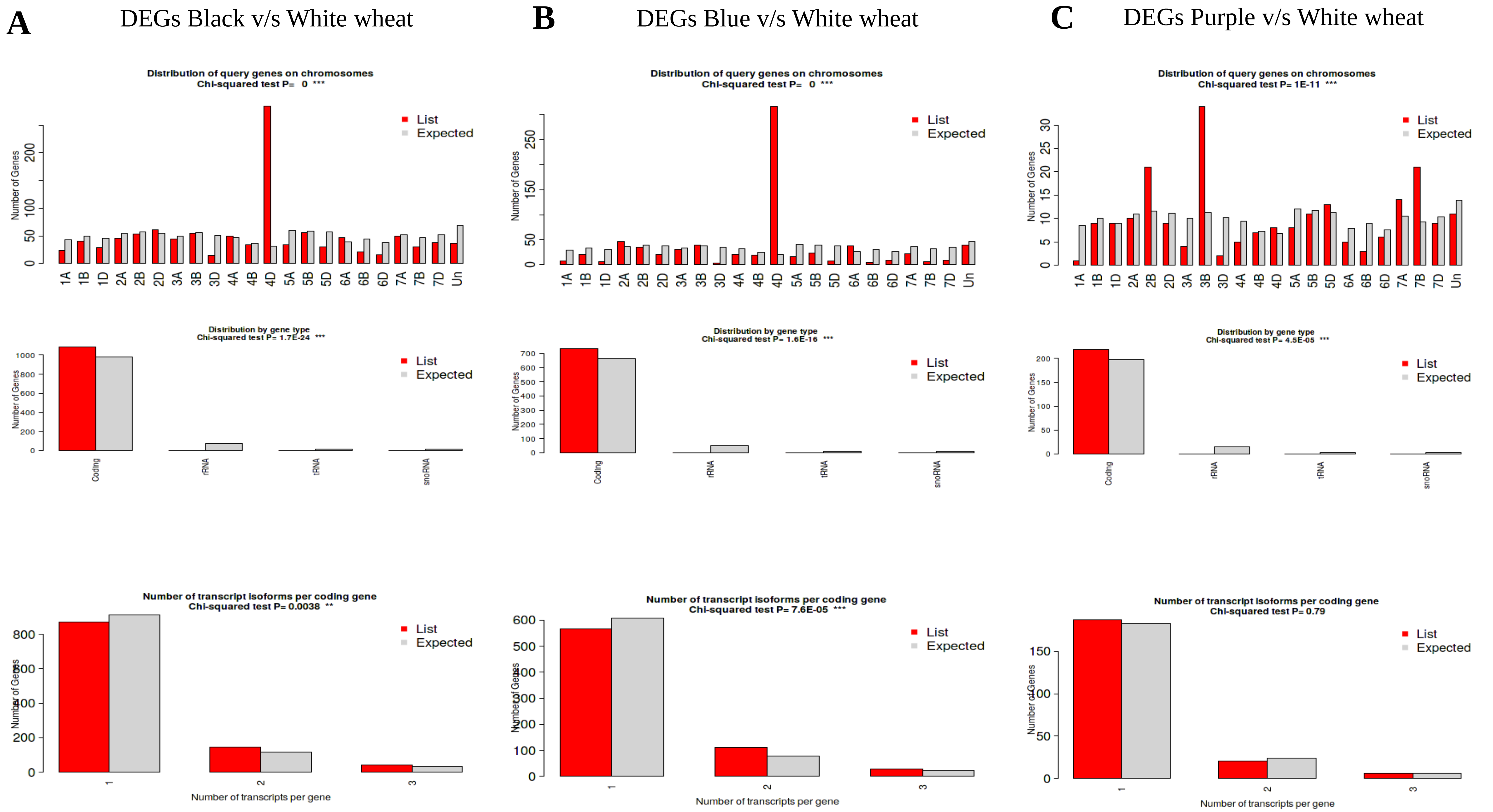


**Figure S2:** Chromosomal distribution, gene type distribution and number of isoforms with chi square test and t-test of differentially expressed genes in **(A)** Black vs white **(B)** Blue vs white **(C)** Purple vs white.

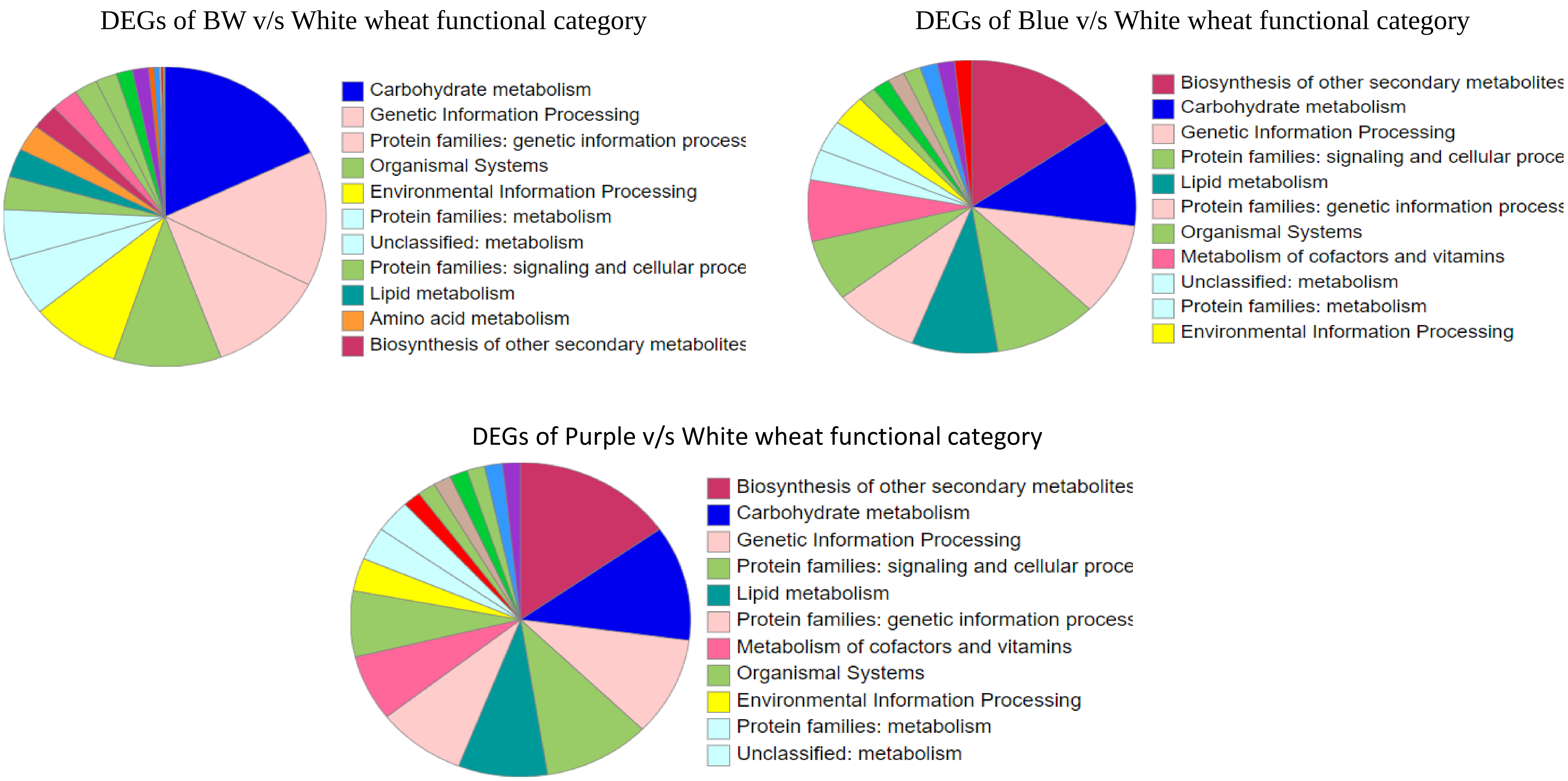


**Figure S3: GO Classification of DEGs of all the three colored wheat samples in Biological Functions**

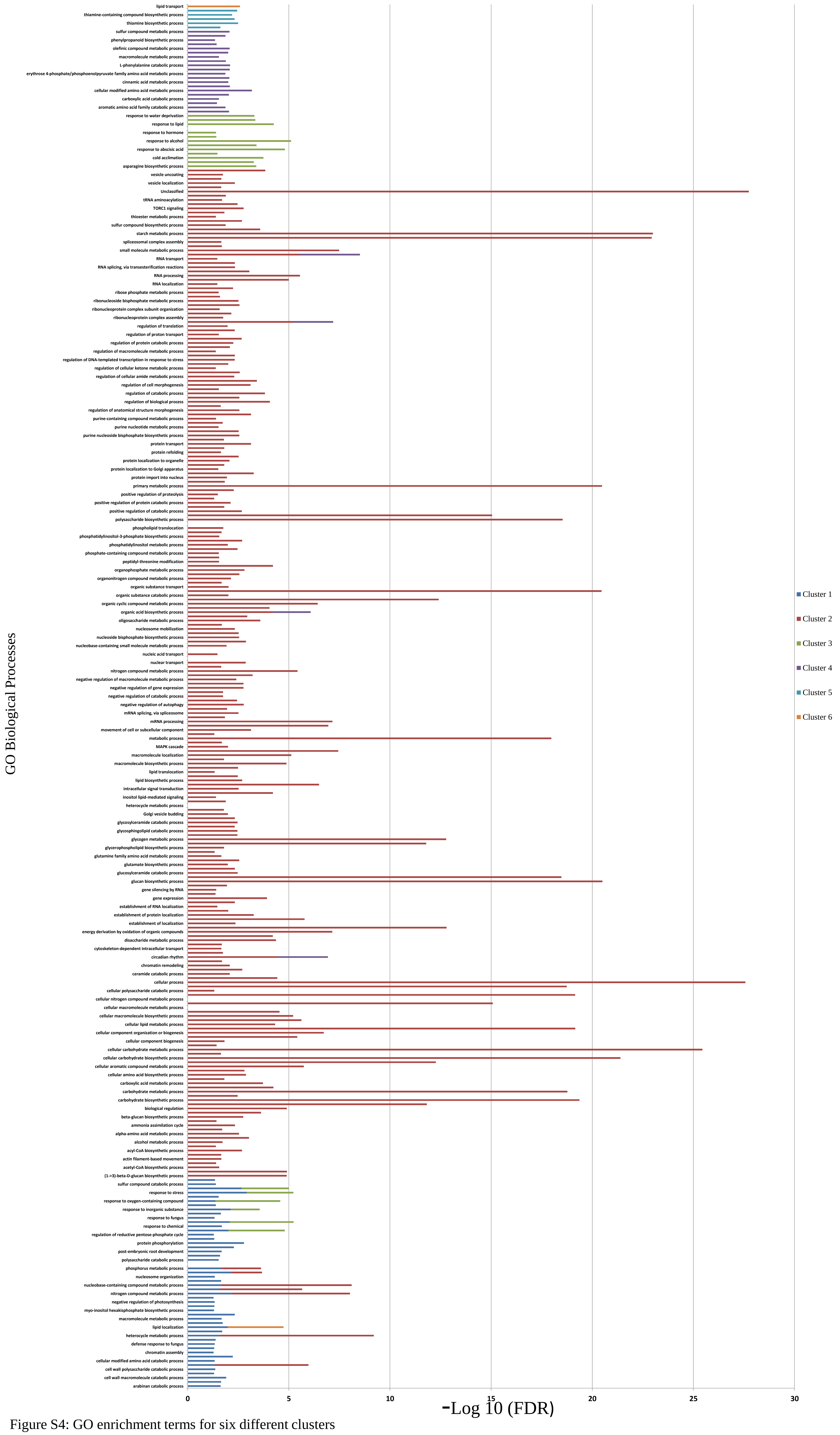


Figure S4: GO enrichment terms for six different clusters

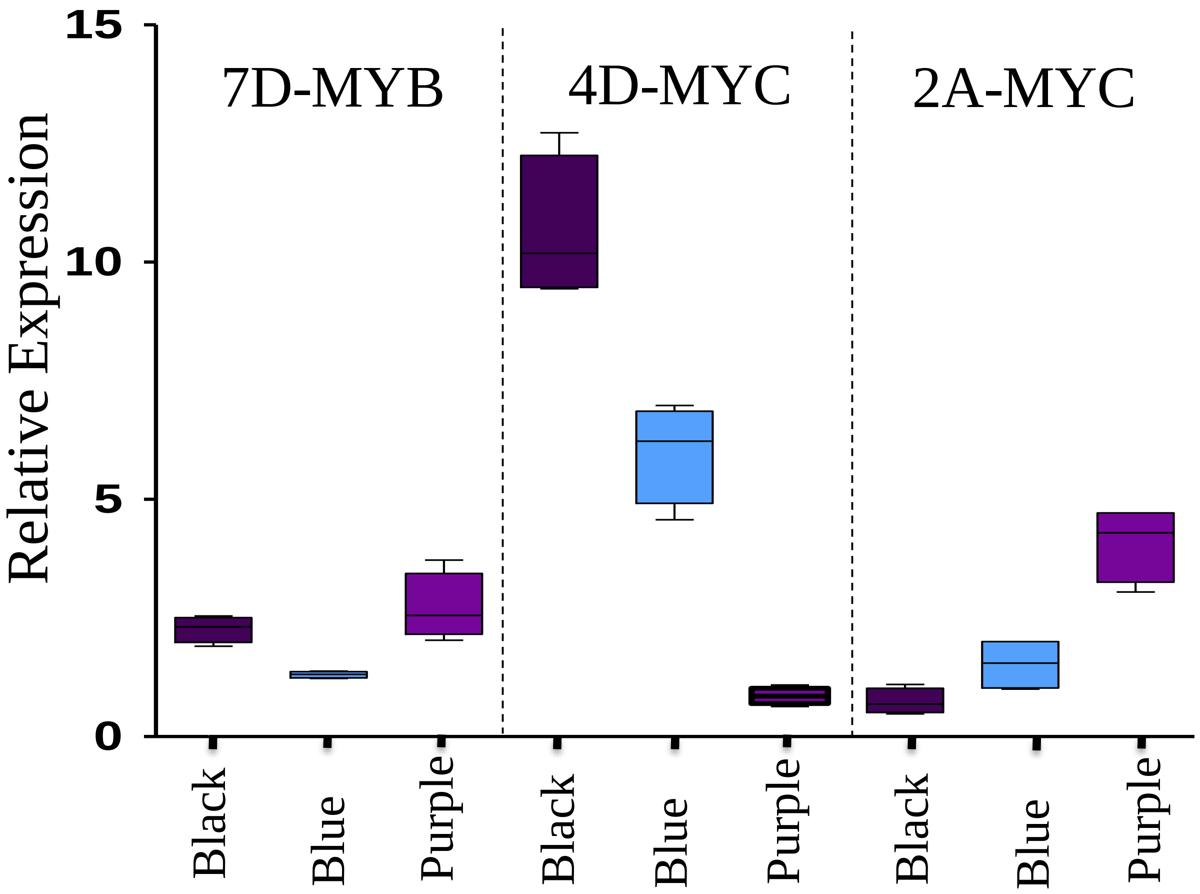


Figure S5: Relative expression of 7D-MYB, 4D-MYC and 2A-MYC in all colored wheat seed at 28DAA /85 Zodax scale stage compared to white wheat.

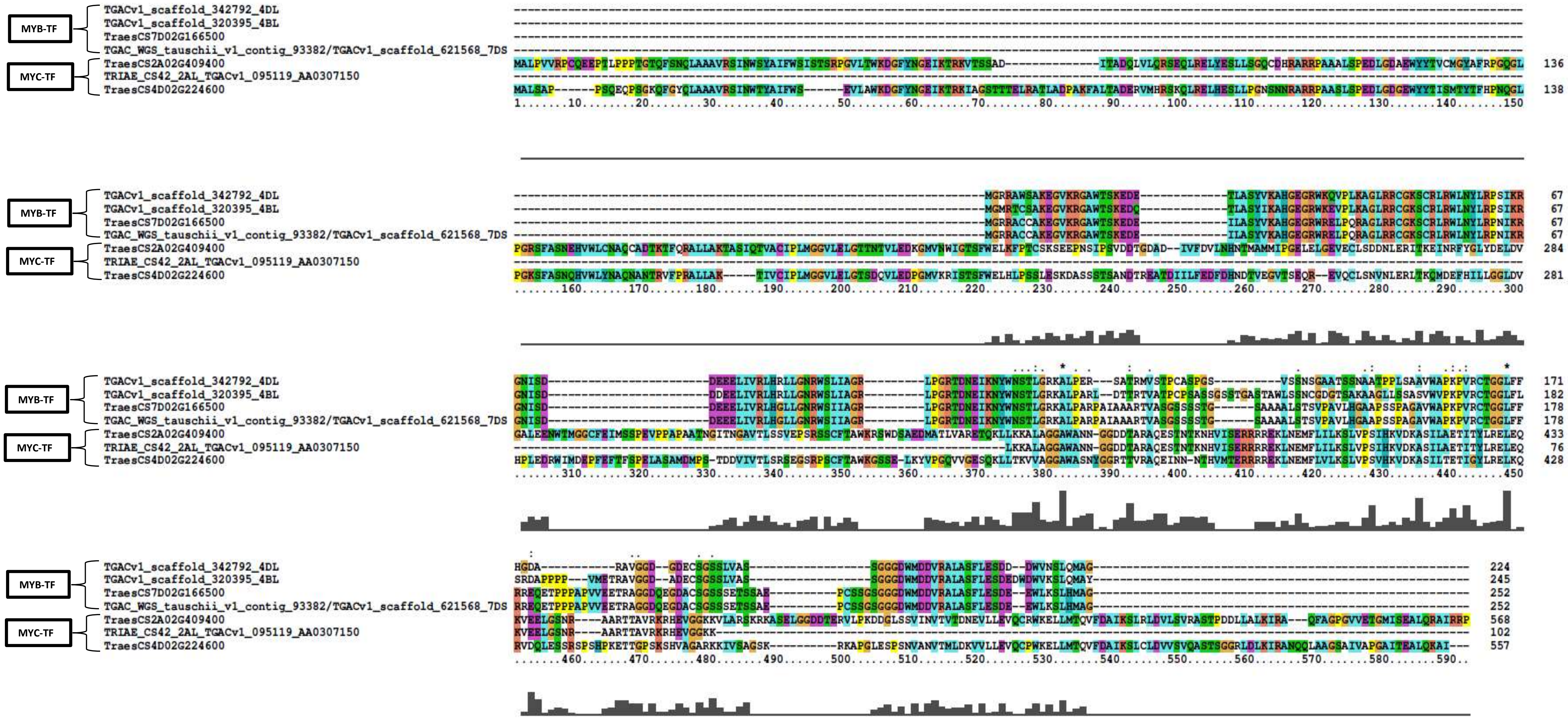


| | |
|---|---|
| TGACv1_scaffold_342792_4DL | TaPpm 2 |
| TGACv1_scaffold_320395_4BL | TaPpm 3 |
| TGAC_WGS_tauschii_v1_contig_93382/TGACv1_scaffold_621568_7DS | TaPpm1 |
| TraesCS7D02G166500 | 7D-myb |
| TRIAE_CS42_2AL_TGACv1_095119_AA0307150 | bHLH motif |
| TraesCS4D02G224600 | 4D-myc |
| TraesCS2A02G409400 | 2A-myc |

Figure S6: Multiple alignments of DE regulatory genes of anthocyanin biosynthesis in our transcriptomics study with reported Pp protein from Jiang *et al*., 2018

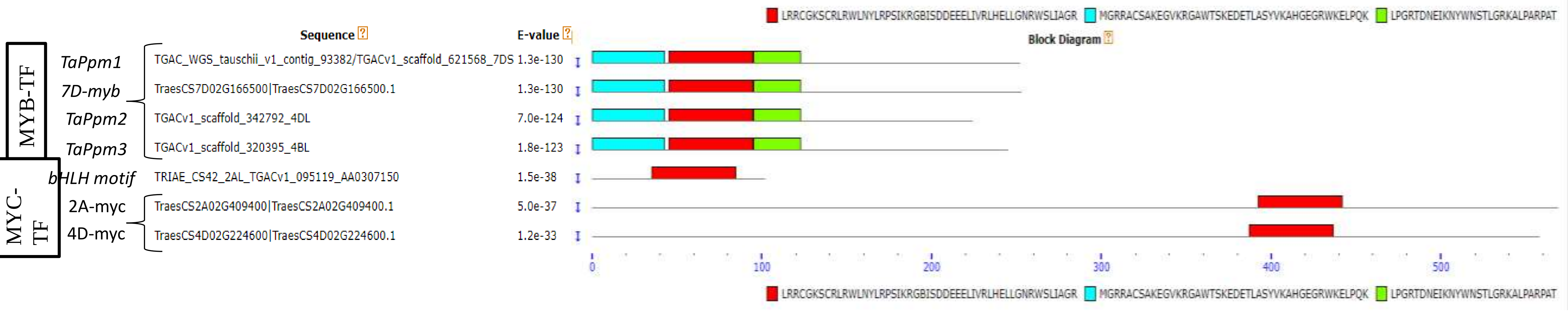


Figure S7: Conserved Motifs of TFs elucidated by MAST.

Colored boxes representing different conserved motifs with different sequences and size.

Where: TaPpm1, TaPpm 2 and TaPpm 3 three MYB-TFs and one bHLH motif sequences were taken from Jiang *et al.*, 2018.
7D-myb, 2A-myc and 4D-myc sequences were taken from own transcriptome analysis results.

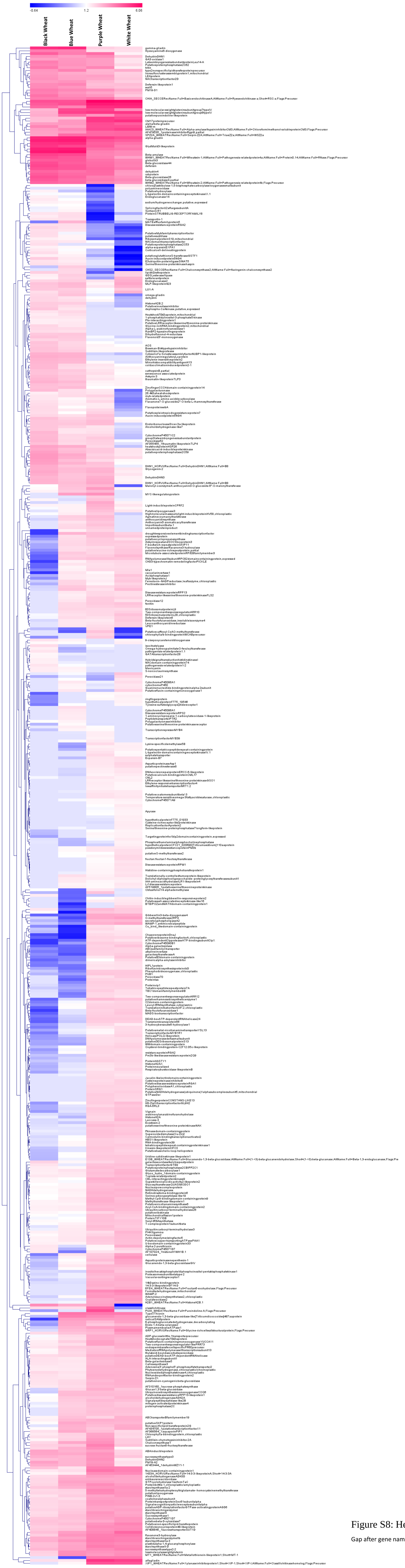


Figure S8: Heat map of all DEGs in three colored wheat seed samples

Gap after gene name depicts the same gene isoform expression pattern

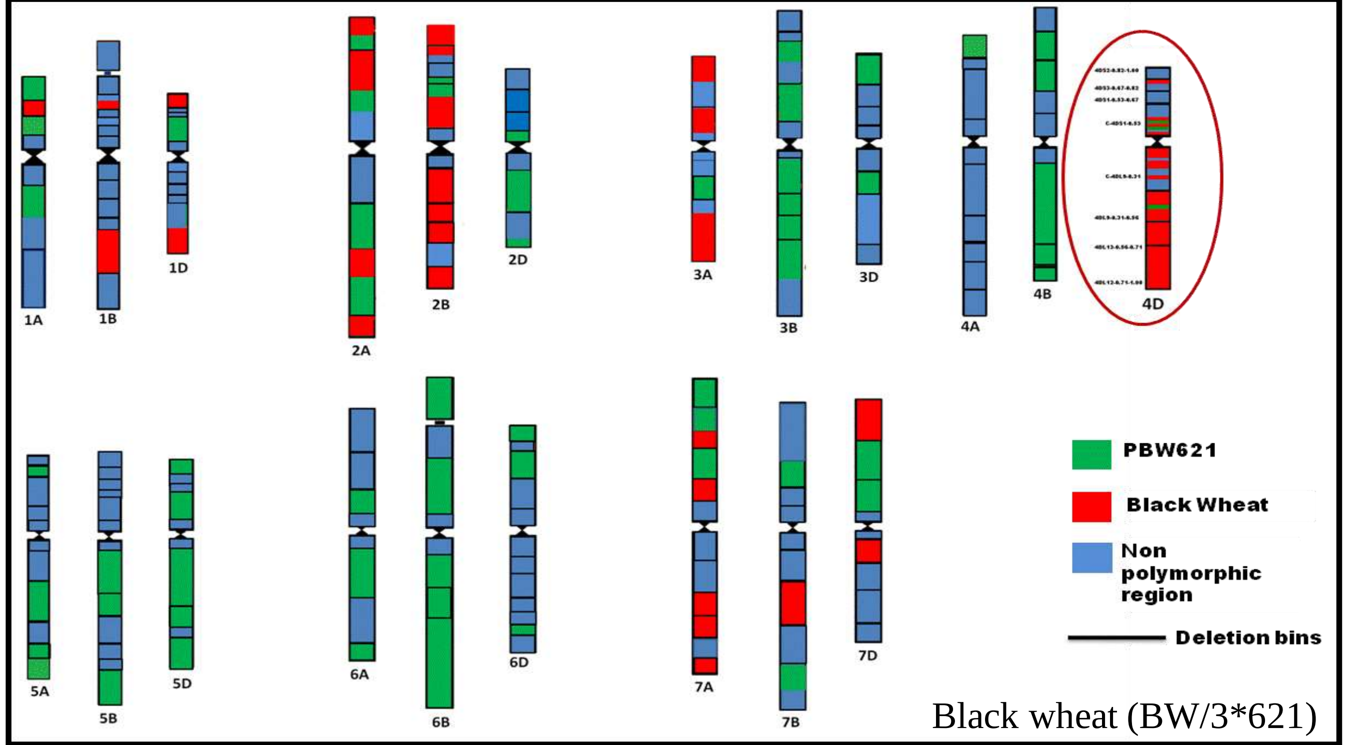


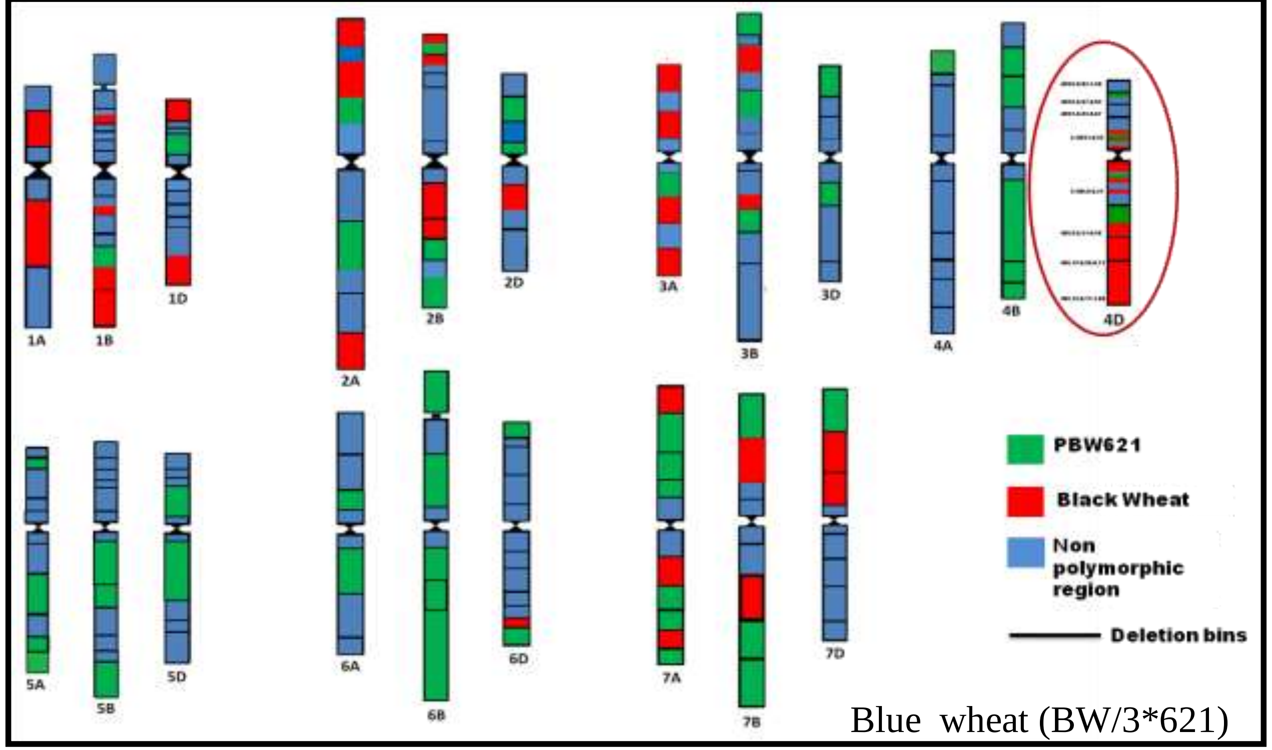


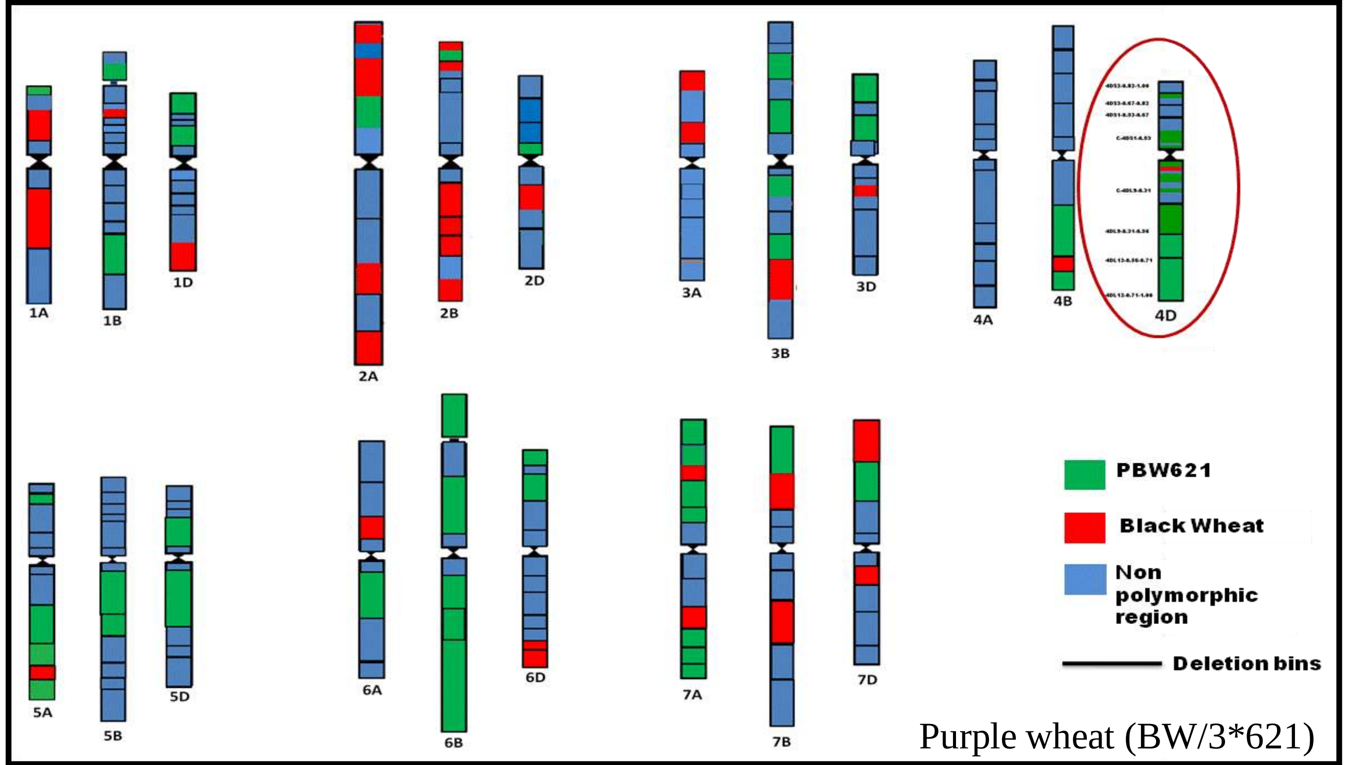


Figure S9: Schematic SSR-based map of black, blue and purple wheat lines segregated from single cross (BW/3*621). Exotci BW donor was created by crossing the blue-colored 4E Agropyron elongatum chromosome substitution line Shou Ien 4E ( 4D) line with a purple colored mutant.

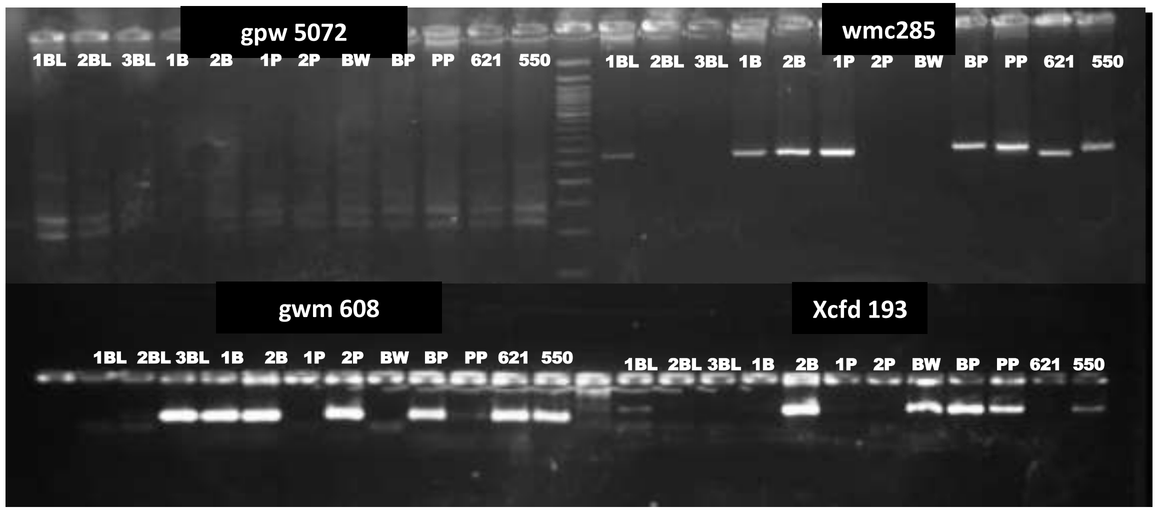


Black wheat (BW/3*621): 1BL-3831, 2BL- 3849, 3BL-3818
Blue wheat(BW/3*621):1B-3921, 2B- 3833
Purple wheat( BW/3*621): 1P- 3840, 2P- 3857
BW: Exotic Black color donor ; BP –TA3972; PP –TA3851; 621, 550 -Cultivars

**Agarose gel (4%) representing polymorphic bands in different color wheat lines along with their parents**

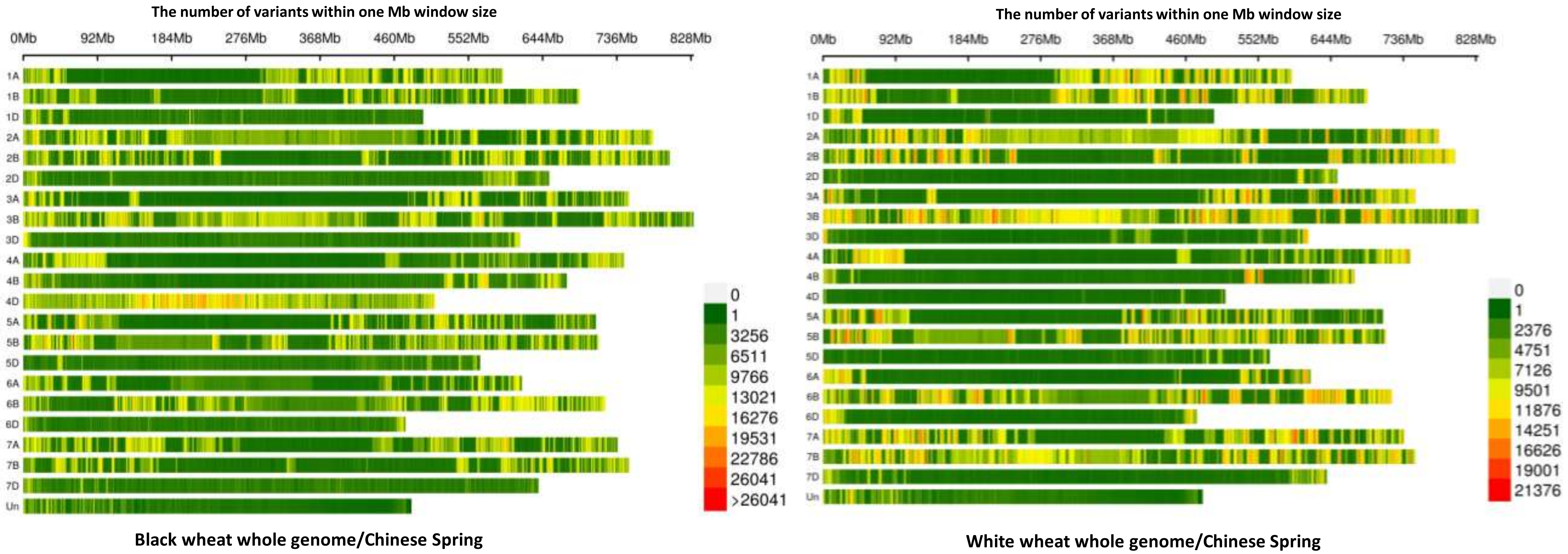


Figure S10**:** Chromosome-wise variations identified in whole genome re-sequenced black and white wheat genotypes in reference to Chinese spring. The variations were identified as SNPs, Insertion, deletion, and complex nucleotide variants.

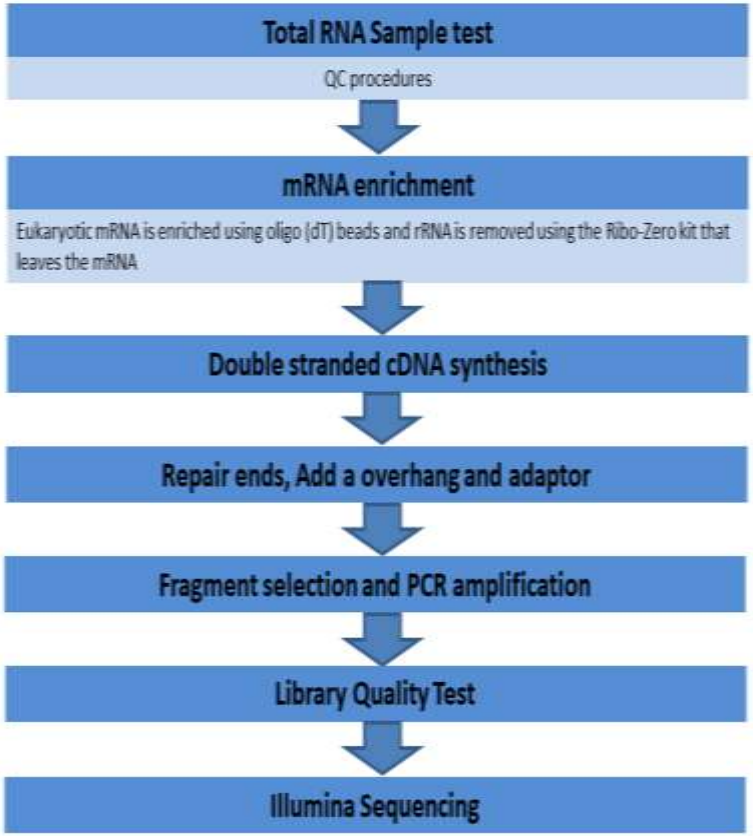


**Figure S11 : Workflow of library construction and RNA sequencing used by outsource company**

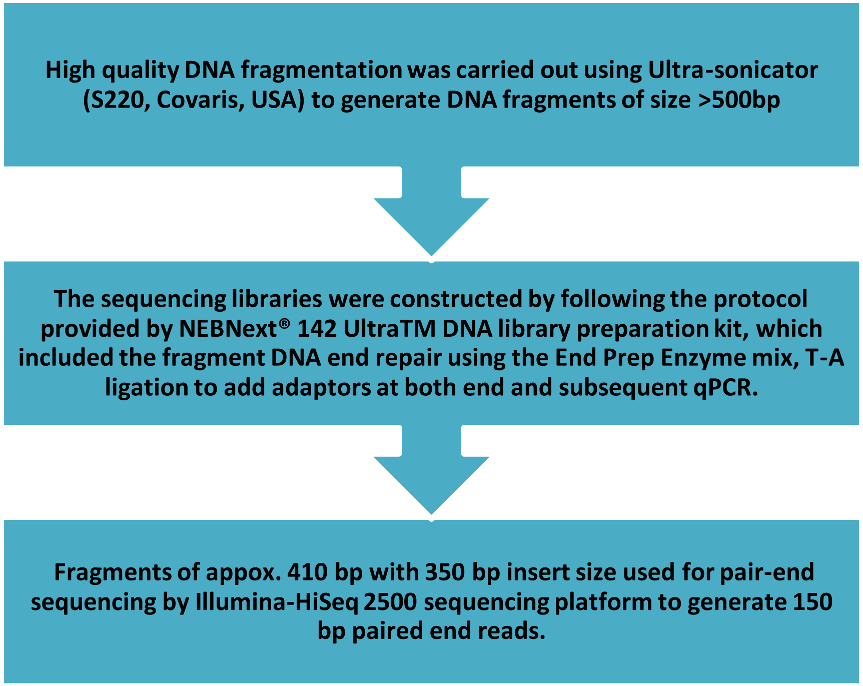


**Figure S12 : Workflow of library construction and DNA sequencing used by outsource company**
