## Supplemental Table 1-4,8,10 for "Transcriptional signature pattern in black, blue and purple wheat and impact on seed pigmentation and other associated features: Comparative transcriptomics, genomics and metabolite profiling"

### Supplementary Data

**Table S1: Total Anthocyanin Content (TAC) in two different environments**

(1) NABI Fields Mohali, Punjab, India (30°44’10” N Latitude at an elevation of 351m above sea level).

(2) Offseason nursery of IIWBR, Keylong, Himachal Pradesh, India (32°30’27.9” N Latitude and 76°59’34’ E Longitude at an elevation of 2971 m above sea level).

| TAC (ppm) | Black Wheat | Blue Wheat | Purple Wheat | PBW621 | Temperature (Day/Night) |
|---|---|---|---|---|---|
| **Punjab** | 140 ±3.1 | 80±2 | 40±2.4 | 5±0.42 | 34.2 ºC/18.6 ºC |
| **Keylong** | 240±2.7 | 120±3.1 | 100±9.8 | 6±1 | 28.6 ºC/16.4 ºC |

ppm: parts per million

#### Table S2: Data quality Summery

| Sample | Raw Reads | Clean Reads | Effective Rate (%) | Q20 (%) | GC Content (%) |
|---|---|---|---|---|---|
| **B1** | 44047963 | 41971165 | 95.29 | 96.89 | 53 |
| **B2** | 43039916 | 41338195 | 96.05 | 96.73 | 52 |
| **B3** | 40606071 | 39083603 | 96.25 | 96.98 | 52 |
| **BL1** | 38915417 | 37650371 | 96.75 | 96.97 | 53 |
| **BL2** | 41057368 | 39948904 | 97.30 | 96.7 | 53 |
| **BL3** | 52853313 | 51484559 | 97.41 | 96.97 | 52 |
| **P1** | 44115644 | 43613212 | 98.86 | 96.84 | 52 |
| **P2** | 52332938 | 51651632 | 98.70 | 96.53 | 52 |
| **P3** | 46580327 | 45057368 | 96.73 | 96. 8 | 52 |
| **W1** | 43039106 | 41825436 | 97.18 | 96.47 | 53 |
| **W2** | 46009667 | 45182985 | 98.20 | 96.72 | 53 |
| **W3** | 54397525 | 52198204 | 95.96 | 96. 7 | 53 |

B1, B2, B3: three replicated of Blue wheat; BL1, BL2, BL3: three replicates of Black wheat; P1, P2, P3: three replicates of Purple wheat; W1, W2, W3: three replicates of White wheat

Where: Effective Rate (%): (Clean reads/Raw reads)*100%;Error rate: base error rate Q20 (Base count of Phred value > 20) / (Total base count); GC content: (G & C base count) / (Total base count)

#### Table S3: Number of Up and Down-regulated DEGs in all colored wheat.

| | Total | Up-regulated | Down-regulated |
|---|---|---|---|
| **Black v/s White** | 1092 | 371 | 721 |
| **Blue v/s White** | 741 | 280 | 461 |
| **Purple v/s White** | 220 | 143 | 77 |

### Table S4: Results of blastP of three putative anthocyanin regulatory TFs with NR database

| Gene | Protein | Name | Accession | Description | Interval | E-value | Top most NCBI BLAST hit with NR-database |
|---|---|---|---|---|---|---|---|
| TraesCS4D02G224600 | Anthocyanin regulatory Lc protein | Name | Accession | Description | Interval | E-value | hypothetical protein CFC21_062292 [Triticum aestivum] |
| | | bHLH-MYC_N | pfam14215 | bHLH-MYC and R2R3-MYB transcription factors N-terminal; This is the N-terminal region of a ... | 20-203 | 3.55E-27 | anthocyanin regulatory R-S protein-like [Aegilops tauschii subsp. strangulata] |
| | | bHLH_AtTT8_like | cd11451 | basic helix-loop-helix (bHLH) domain found in Arabidopsis thaliana protein transparent testa 8 ... | 379-440 | 2.67E-22 | MYC4E [Thinopyrum obtusiflorum] |
| | | HLH | smart00353 | helix loop helix domain; | 386-432 | 4.51E-14 | MYC & bHLH Hordeum Vulgure |
| | | HLH | pfam00010 | Helix-loop-helix DNA-binding domain; | | | anthocyanin regulatory R-S protein isoform X2 [Aegilops tauschii subsp. strangulata] |

| Gene | Protein | Name | Accession | Description | Interval | E-value | Top most NCBI BLAST hit with NR-database |
|---|---|---|---|---|---|---|---|
| TraesCS2A02G409400 | MYC-like regulatory protein | Name | Accession | Description | Interval | E-value | MYC-like regulatory protein [Triticum aestivum] |
| | | bHLH-MYC_N | pfam14215 | bHLH-MYC and R2R3-MYB transcription factors N-terminal; This is the N-terminal region of a ... | 25-206 | 2.47E-39 | MYC1 isoform III [Triticum aestivum] |
| | | bHLH_AtTT8_like | cd11451 | basic helix-loop-helix (bHLH) domain found in Arabidopsis thaliana protein transparent testa 8 ... | 384-452 | 2.01E-21 | bHLH transcription factor [Triticum aestivum] |
| | | HLH | smart00353 | helix loop helix domain; | 389-437 | 1.82E-14 | anthocyanin regulatory R-S protein-like isoform X1 [Triticum dicoccoides] |
| | | HLH | pfam00010 | Helix-loop-helix DNA-binding domain; | | | |

| Gene | Protein | Name | Accession | Description | Interval | E-value | Top most NCBI BLAST hit with NR-database |
|---|---|---|---|---|---|---|---|
| TraesCS7D02G166500 | myb-related protein | Name | Accession | Description | Interval | E-value | anthocyanin regulatory C1 protein [Aegilops tauschii subsp. strangulata] |
| | | PLN03212 | PLN03212 | Transcription repressor MYB5; Provisional | 6-115 | 1.26E-59 | MYB-A1 protein [Triticum monococcum] |
| | | REB1 | COG5147 | Myb superfamily proteins, including transcription factors and mRNA splicing factors ... | 12-114 | 1.68E-19 | myb-related protein A1 [Triticum aestivum] |
| | | Myb_DNA-binding | pfam00249 | Myb-like DNA-binding domain; This family contains the DNA binding domains from Myb proteins, ... | 67-112 | 6.06E-13 | MYB-A1 protein [Triticum monococcum] |
| | | SANT | smart00717 | SANT SWI3, ADA2, N-CoR and TFIIIB'' DNA-binding domains; | 14-63 | 1.17E-09 | |
| | | SANT | cd00167 | 'SWI3, ADA2, N-CoR and TFIIIB' DNA-binding domains. Tandem copies of the domain bind telomeric ... | | | |

**Table S8 :** Primers used for background screening with their position in chromosome

| S.No. | Primers | Position | S.No. | Primers | Position | S.No. | Primers | Position | S.No. | Primers | Position | S.No. | Primers | Position | S.No. | Primers | Position |
|---|---|---|---|---|---|---|---|---|---|---|---|---|---|---|---|---|---|
| 1 | gpw2186 | 1AS | 29 | wmc153 | 1DL | 57 | gpw5237 | 2DL | 85 | gpw1149 | 3DL | 114 | gpw7574 | 5BL | 142 | gwm325 | 6DS |
| 2 | wmc24 | 1AS | 30 | wmc382 | 2AS | 58 | gpw5239 | 2DL | 86 | gpw5235 | 3DL | 115 | gpw3191 | 5BL | 143 | gpw5076 | 6DL |
| 3 | gpw7006 | 1AS | 31 | gpw2018 | 2AS | 59 | Cfd 2 | 3AS | 87 | gpw2239 | 4AS | 116 | gwm408 | 5BL | 144 | gpw7303 | 6DL |
| 4 | gpw2277 | 1AS | 32 | gpw7399 | 2AS | 60 | gpw4074 | 3AS | 88 | wmc173 | 4AS | 117 | gpw3183 | 5BL | 145 | gpw7288 | 7AS |
| 5 | gpw4331 | 1AL | 33 | gpw3167 | 2AS | 61 | gpw4221 | 3AS | 89 | gpw1010 | 4AS | 118 | wmc233 | 5DS | 146 | gpw8289 | 7AS |
| 6 | gpw7258 | 1AL | 34 | gpw3249 | 2AL | 62 | gpw3036 | 3AS | 90 | gpw7253 | 4AL | 119 | gpw4467 | 5DS | 147 | gpw3140 | 7AS |
| 7 | gpw4410 | 1AL | 35 | gwm312 | 2AL | 63 | gpw4401 | 3AL | 91 | gpw3224 | 4AL | 120 | gpw4475 | 5DS | 148 | wmc603 | 7AS |
| 8 | gpw4373 | 1AL | 36 | gpw2206 | 2AL | 64 | gpw2266 | 3AL | 92 | gpw4120 | 4BS | 121 | gpw5129 | 5DL | 149 | gpw7185 | 7AS |
| 9 | gpw2067 | 1BS | 37 | gpw8034 | 2AL | 65 | gpw5271 | 3AL | 93 | gpw3125 | 4BS | 122 | cfd8 | 5DL | 150 | gpw3038 | 7AL |
| 10 | wmc329 | 1BS | 38 | gpw4218 | 2BS | 66 | gpw5175 | 3AL | 94 | gpw4101 | 4BL | 123 | gwm271 | 5DL | 151 | wmc525 | 7AL |
| 11 | gpw4069 | 1BS | 39 | wmc764 | 2BS | 67 | gpw8020 | 3BS | 95 | gpw4410 | 4BL | 124 | cfd183 | 5DL | 152 | gpw3276 | 7AL |
| 12 | wmc128 | 1BS | 40 | wmc157 | 2BS | 68 | wmc78 | 3BS | 96 | gwm495 | 4BL | 125 | gpw7107 | 5DL | 153 | gpw5180 | 7AL |
| 13 | gpw3044 | 1BS | 41 | wmc83 | 2BS | 69 | wmc231 | 3BS | 97 | gpw4175 | 4BL | 126 | gpw7592 | 6AS | 154 | gpw3127 | 7AL |
| 14 | gpw7422 | 1BS | 42 | gpw1109 | 2BS | 70 | wmc777 | 3BS | 98 | gpw4189 | 4DS | 127 | gpw4347 | 6AS | 155 | gpw4100 | 7AL |
| 15 | gpw3010 | 1BS | 43 | gpw4382 | 2BS | 71 | gwm274 | 3BL | 99 | gpw5072 | 4DS | 128 | gpw4344 | 6A CENTROMERE | 156 | gpw4101 | 7BS |
| 16 | gpw7812 | 1B CENTROMERE | 44 | gpw3032 | 2BL | 72 | gpw1145 | 3BL | 101 | wmc720 | 4DL | 129 | gpw2222 | 6AL | 157 | gpw4444 | 7BL |
| 17 | gpw4478 | 1BL | 45 | gpw3090 | 2BL | 73 | gpw4331 | 3BL | 102 | gpw4538 | 4DL | 130 | wmc256 | 6AL | 158 | gpw5058 | 7BL |
| 18 | gwm498 | 1BL | 46 | gpw4103 | 2BL | 74 | gpw1025 | 3BL | 103 | gpw5255 | 4DL | 131 | gpw7388 | 6AL | 159 | gpw7596 | 7BL |
| 19 | gpw7603 | 1BL | 47 | gpw7506 | 2BL | 75 | gpw5016 | 3BL | 104 | gpw7607 | 5AS | 132 | gpw4357 | 6BS | 160 | gpw4471 | 7BL |
| 20 | wmc156 | 1BL | 48 | gpw1214 | 2BL | 76 | gpw7335 | 3BL | 105 | gpw7218 | 5AS | 133 | gpw4032 | 6BS | 161 | gpw8090 | 7BL |
| 21 | gpw7646 | 1BL | 49 | gpw4124 | 2DS | 77 | gpw4044 | 3BL | 106 | gpw3003 | 5AS | 134 | wmc95 | 6BS | 162 | gpw5208 | 7DS |
| 22 | gpw5162 | 1BL | 50 | gpw5155 | 2DS | 78 | gpw7321 | 3DS | 107 | gpw4228 | 5AL | 135 | gpw5212 | 6BS | 163 | gpw299 | 7DS |
| 23 | gpw1077 | 1BL | 51 | gpw5176 | 2DS | 79 | gpw333 | 3DS | 108 | gpw2059 | 5AL | 136 | gpw3122 | 6BL | 164 | gpw4358 | 7DS |
| 24 | gpw7443 | 1BL | 52 | gpw5212 | 2DS | 80 | gpw5104 | 3DS | 109 | gpw4457 | 5AL | 137 | gpw3241 | 6BL | 165 | gpw5211 | 7DS |
| 25 | wmc147 | 1DS | 53 | gpw7325 | 2DS | 81 | gpw4101 | 4BS | 110 | gpw2136 | 5AL | 138 | gpw5076 | 6BL | 166 | gpw5037 | 7DL |
| 26 | wmc329 | 1DS | 54 | gpw7285 | 2DL | 82 | gwm497 | 3DS | 111 | gwm291 | 5AL | 139 | gpw7303 | 6BL | | | |
| 27 | gwm337 | 1DS | 55 | gpw7285 | 2DL | 83 | gpw5166 | 3D CENTROMERE | 112 | gpw7180 | 5BS | 140 | gpw4357 | 6DS | | | |
| 28 | gpw2118 | 1DL | 56 | gpw1247 | 2DL | 84 | gpw7467 | 3DL | 113 | gpw4362 | 5BL | 141 | gpw7079 | 6DS | | | |

**Table S10 :** 4D specific Primers with their position

| S.No. | Primers | Position | S.No. | Primers | Position | S.No. | Primers | Position |
|---|---|---|---|---|---|---|---|---|
| 1. | Xwmc285 | 4DS | 13. | Xcfd23 | 4DL | 25. | Xcfd84 | 4DL |
| 2. | Xwmc818 | 4DS | 14. | Xwmc473 | 4DL | 26. | Xwmc622 | 4DL |
| 3. | Xwmc720 | 4DS | 15. | Xgwm133 | 4DL | 27. | Xgpw5255 | 4DL |
| 4. | Xgpw2220 | 4DS | 16. | Xwmc182 | 4DL | 28. | Xgwm194 | 4DL |
| 5. | Xwmc48 | 4DS | 17. | Xwmc489 | 4DL | 29. | Xwmc825 | 4DL |
| 6. | Xgpw4189 | 4DS | 18. | Xwmc457 | 4DL | 30. | Xgwm624 | 4DL |
| 7. | Xcfd106 | 4DS | 19. | Xgwm165 | 4DL | 31. | Xgwm609 | 4DL |
| 8. | Xwmc52 | 4DS | 20. | Xwmc33 | 4DL | 32. | gpw4189 | 4DS |
| 9. | Xgwm213 | 4DS | 21. | Xwmc331 | 4DL | 33.. | gpw5072 | 4DS |
| 10. | Xgwm608 | 4DS | 22. | Xwmc206 | 4DL | 34.. | wmc720 | 4DL |
| 11. | Xbarc98 | 4DS CENTROMERE | 23. | Xwmc399 | 4DL | 35. | gpw4538 | 4DL |
| 12. | Xcfd193 | 4DS CENTROMERE | 24. | Xcfd39 | 4DL | 36. | gpw5255 | 4DL |
